# *In vivo* optimized three-photon imaging of intact mouse tibia links plasma cell motility to functional states in the bone marrow

**DOI:** 10.1101/2023.12.16.571998

**Authors:** Asylkhan Rakhymzhan, Alexander F. Fiedler, Robert Günther, Scott Domingue, Laura Wooldridge, Ruth Leben, Yu Cao, Anne Bias, Jay Roodselaar, Ralf Köhler, Carolin Ulbricht, Judith Heidelin, Volker Andresen, Ingeborg Beckers, Astrid Haibel, Georg Duda, Anja E. Hauser, Raluca A. Niesner

**Affiliations:** German Rheumatism Research Center – a Leibniz Institute, Biophysical Analytics, Berlin, Germany; German Rheumatism Research Center – a Leibniz Institute, Immune Dynamics, Berlin, Germany; Charité – Universitätsmedizin, Berlin, corporate member of Freie Universität Berlin and Humboldt-Universität zu Berlin, Clinics for Rheumatology and Clinical Immunology, Berlin, Germany; Freie Universität Berlin, Dynamic and Functional in vivo Imaging, Berlin, Germany; Thorlabs Inc., Colorado, US; Medical Physics – Berlin School of Applied Sciences, Berlin, Germany; Miltenyi Biotec GmbH, Bielefeld site, Germany; Charité – Universitätsmedizin, Berlin, corporate member of Freie Universität Berlin and Humboldt-Universität zu Berlin, Julius Wolff Institute, Berlin, Germany

**Author notes:** A.R. and A.F.F. equally contributing first authors. R.A.N. and A.E.H. equally supervised the work.

## Abstract

Intravital multi-photon imaging of the bone marrow is crucial to the study of cellular dynamics, communication with the microenvironment and functions, however, imaging of deep tissue areas is challenging and minimally invasive methods for deep-marrow imaging in intact long bones are needed. We developed a high pulse energy 1650 nm laser prototype, which permits to surpass >100 µm thick cortical bone and to perform three-photon microscopy (3PM) in more than 400 µm depth in the marrow cavity of intact mouse tibia *in vivo*. Its unique 3 and 4 MHz laser repetition rates allowed us to analyze motility patterns of rare cells over large fields of view deep within the unperturbed marrow. In this way, we found a bi-modal migratory behavior of marrow plasma cells. Besides, the analysis of third harmonics generation (THG) in the tibia identified this signal to be a label-free indicator of the abundance of cellular organelles, in particular the endoplasmic reticulum, reflecting protein biosynthesis capacity. We found that only one third of the plasma cells in the tibia marrow of adult mice have a strong THG signal and, thus, a high protein synthesis capacity, while the other two thirds of plasma cells display a low THG signal. Finally, we identified an inverse link between migratory behavior and THG signal strength in marrow plasma cells. As in these cells, the protein biosynthesis capacity indicated by a strong THG signal is mainly associated with antibody secretion, we could relate motility to functional states of plasma cells *in vivo*. Our 3PM method retains the ability to connect cellular dynamics to protein biosynthesis capacity in various marrow cell types beyond plasma cells, as THG is a ubiquitous signal, opening new perspectives on understanding how tissue microenvironment impacts on cellular functions in the bone marrow.

## Introduction

Intravital two-photon microscopy (2PM) in flat^1^ and long bones of mice^2,3^ allows to study cellular migration, communication and functions in bone marrow. As a primary lymphoid organ, the bone marrow is the birthplace of immune cells. This includes the generation of B cells, however, it also constitutes the final destination for terminally differentiated plasma cells, emerging from B cells activated in the periphery. Marrow plasma cells ensure long-term immunological memory through antibody secretion^4^. *In vivo* 2PM has helped us to better understand plasma cell dynamics and interactions in the bone marrow microenvironment, on various time scales^5–7^, which is essential for their long-term survival *in situ*^8^. Besides, *in vivo* bone marrow imaging in both flat and long bones has brought us insights into the development and dynamics of hematopoietic stem cells^9^, osteoclast migration^10,11^, as well as into tumor cell dormancy^12^ and metastasis dissemination^13,14^. In long bones, *in vivo* 2PM has informed us about dynamic changes within mesenchymal and vascular compartments, in bone and marrow tissue^15,16^, revealing the broad utility of this technology.

Recently, distinct molecular fingerprints and functions of the marrow tissue in the skull have been demonstrated^17^, different from those of the marrow tissue in long bones. This finding emphasizes the need for dedicated *in vivo* imaging technologies for each bone type.

Although optical imaging using near-infrared 2PM (700-1350 nm, ≈100 MHz repetition rate, pulse energy < 5 nJ) in lymph nodes^18,19^, spleen^20^ and hematopoietic islets in intact flat bones, e.g. calvarium^21^, is broadly applied, scattering and absorption of radiation next to wave-front distortions impair the image quality with increasing tissue depth. *In vivo* near-infrared 2PM in the marrow cavity of long bones through intact bone cortex has been successfully performed^10,11^, however, only in mice with thin bone cortex, such as young or γ-irradiated mice, and at shallow tissue depths^22^. *In vivo* 2PM in long bones of adult mice, with thick bone cortex, has been previously performed either by mechanically thinning the overlaying bone tissue^5,15,23^, or by inserting fixed micro-endoscopic probes in the femur^24^. These approaches, applicable in all mice independent of their age, require invasive bone surgery, which initially causes an immune reaction and, by that, a transient perturbation of the tissue environment at the imaging site^24^. Bone has the highest refractive index span among mammalian tissues (from n ≈1.33 up to ≈1.62)^25–27^. For comparison, the refractive index in soft lymphoid tissues ranges between ≈1.33 and ≈1.4^26–28^. Therefore, perturbation-free *in vivo* imaging of the marrow cavity through intact thick bone cortex in long bones cannot be achieved using near-infrared 2PM, due to strong scattering effects in calcified bone.

In response to the challenges in near-infrared 2PM, three-photon microscopy (3PM) using high pulse energy infrared radiation has been developed, to allow access to deeper tissue layers in various organs^29–32^. Technological improvements such as the design of 3PM with Bessel beam illumination^33,34^, multicolor 3PM^35^, adaptive-optics enhanced 3PM^36,37^, or miniaturized 3PM in freely moving mice^38^ helped to perform imaging at unprecedented tissue depths in brain cortex and hippocampus. Moreover, 3PM allowed for structural and functional murine brain imaging through the intact skull (up to 100 µm thick)^32^, which requires wave-front distortion correction using customized adaptive optics^36^. Besides, deep-tissue *in vivo* intra-tumoral imaging in skin and *ex vivo* imaging of ectopic ossicles with ≈70 µm thick cortical bone have been demonstrated^39^. Next to fluorescence detection, 3PM enables the detection of third harmonics generation (THG), an elastic three-photon scattering process of coherent radiation, which provides label-free information about periodically organized molecular structures and about refractive index heterogeneity, e.g. in lipid bi-layers of cell or organelle membranes. In mouse tissue, THG has been associated with both cellular and extracellular structures^40^ and has been used to study myelination in the brain cortex^41^, the geometry of osteocytes in the bone matrix^42^ or blood flow^29^, relying on the THG signal of erythrocytes. Besides, oxygenation has recently been demonstrated to be indicated by THG and sum frequency generation, when using two spatiotemporally synchronized laser pulse trains^43^. In leukocytes, a strong THG signal was shown to correlate with high cellular granularity^44^.

Addressing the need to monitor immune cell migration in deep regions of lymphoid organs^45^, a customized 3PM setup has been designed to enable *in vivo* imaging throughout murine naïve lymph nodes and in spleens. Hence, lymphocyte dynamics could be analyzed down to ≈600 µm depth in these secondary lymphoid organs^29^. Although being a promising method for dynamic *in vivo* deep-marrow imaging in intact long bones of adult mice, high pulse energy infrared 3PM has neither been used in, nor specifically adapted to this type of application.

Laser systems adequate as excitation sources^31,35^ for successful deep-tissue 3PM must:

i. emit at a radiation wavelength in one of the infrared spectral windows: 1300-1400 nm and 1600-1700 nm, respectively, for reduced signal loss in tissue,
ii. have a high pulse energy and low repetition rate to secure moderate, biocompatible average power, and
iii. feature temporally narrow pulses (≈100 fs), to provide high photon density at the sample for efficient three-photon excitation.

Based on pulse chirping to support the generation of high pulse energy between 100 nJ and few µJ, different configurations of optical parametric amplifiers (OPA) have been used as excitation sources for three-photon imaging in living tissue^29,31,35^. Typically, OPA lasers are tunable between 1200 and 1700 nm at repetition rates from 0.33 to 1 MHz (rarely 2 MHz). The low repetition rate represents a disadvantage for *in vivo* 3PM, as it limits the image acquisition speed over fields of view spanning several hundreds of µm. Soliton lasers have been successfully used as excitation sources to increase imaging depth in the mouse brain cortex by *in vivo* 2PM, owing to their tunable repetition rates up to 10 MHz^46,47^. The high repetition rates are favorable for increased image acquisition speed. However, their moderate pulse energy (<50 nJ) is potentially too low to efficiently induce three-photon events in tissue.

Here, we developed and integrated a novel OPA prototype in a customized multi-photon microscope system, to enable dynamic *in vivo* three-photon imaging in the marrow cavity of the intact tibia in adult mice. The high pulse energy radiation at 1650 nm enabled us to image marrow tissue at a subcellular resolution through up to 200 µm thick bone cortex, reaching imaging depths larger than 400 µm in the tibial marrow. By tuning the OPA repetition rate up to 4 MHz, we were additionally able to conduct time-lapse 3PM over large fields of view (400×400 µm²) in the deep marrow cavity of intact unperturbed tibia *in vivo*. In this way, we visualized blood flow in the bone marrow vasculature and analyzed the migration of even rare cell subsets, such as plasma cells. Taking advantage of label-free THG signals, we found that plasma cells’ functional state and migration behavior *in vivo* are tightly linked.

## Results

### Microscope design for dynamic *in vivo* deep-tissue imaging in long bones

*In vivo* imaging of intact long bones requires the excitation laser beam to surpass two types of tissue layers: the hard cortical bone and the bone marrow, a soft lymphoid tissue. Laser power attenuation in both tissue types is caused by scattering and absorption of radiation, in a wavelength-dependent manner^28,48^. In the infrared spectral range, the absorption of radiation displays local minima at 1300 nm and 1650 nm, being lower in bone than in soft lymphoid tissue, due to the high water content of the latter^27,28,49^. Notably, the absorption is stronger at 1650 nm than at 1300 nm for both tissue types. In contrast, the scattering of radiation is stronger in bone than in lymphoid tissue at a given wavelength, and it steadily decreases with increasing wavelength^27^. Taken together, the signal attenuation in bone tissue is lower at 1650 nm (8.7 cm^-1^) compared to 1350 nm (9.9 cm^-1^), and generally higher than in lymphoid tissues (5.4 cm^-1^ at 1330 nm and 7.1 cm^-1^ at 1650 nm). Calculations were made based on a previously published model of radiation scattering^28^ and absorption spectra^27^ for both tissue types. Hence, we expected a superior performance of 1650 nm compared to 1350 nm excitation for deep-tissue *in vivo* imaging through thick (> 100 µm) tibia cortex.

To test this assumption and to retrieve adequate experimental parameters for fast dynamic deep-tissue tibia imaging over large 3D volumes, we adapted and characterized a state-of-the-art two-photon microscope setup to enable efficient two- and three-photon imaging in a broad infrared wavelength range (Fig. 1, Suppl. Fig. 1, **Table I**), as described in detail in *Methods* and *Suppl. Info*. Besides lasers typically used for two-photon microscopy (i.e. Ti:Sa and an optical parametric oscillator (OPO) operating at 80 MHz repetition rate in the range 700-1080 nm and 1050-1350 nm, respectively), we implemented a novel optical parametric amplifier (OPA) as an excitation source. This OPA features a fixed wavelength (1650 nm, spectral bandwidth Δλ = 60 nm) and variable repetition rates (1.01-3.98 MHz), at fixed pulse energy and 65 fs pulse width (**Table I**, Suppl. Fig. 1). The novel laser is termed hereafter Ytterbia OPA. For comparison, we used a state-of-the-art OPA laser, tunable in the range 1200 nm – 1700 nm, at a fixed repetition rate of 2 MHz and < 60 fs temporal pulse width.

**Figure 1.**
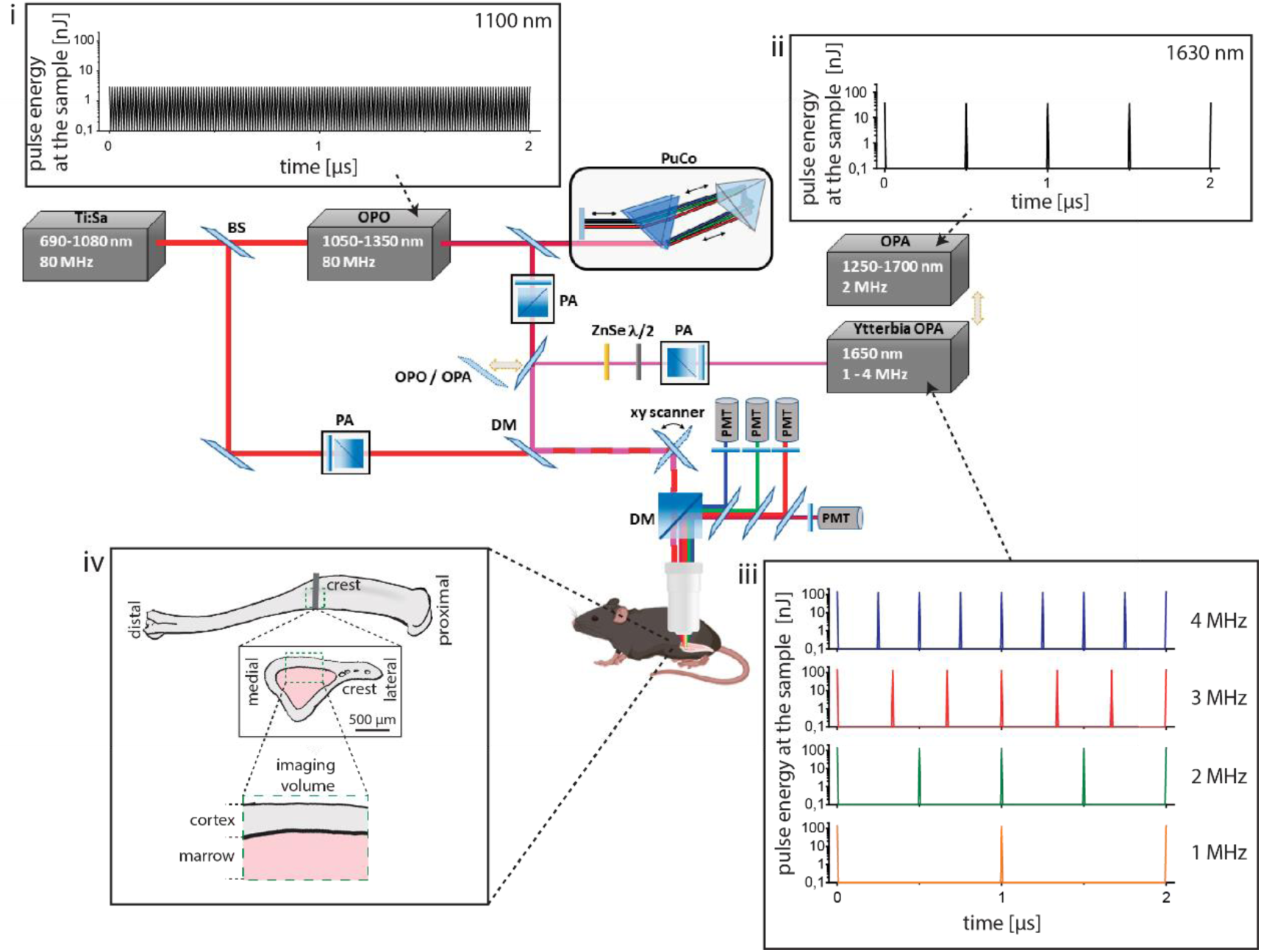
Customized setup design for dynamic deep-tissue imaging within intact long bones *in vivo*. The schematics depicts the main components of the customized multi-photon microscope. State-of-the-art two-photon imaging is performed using low pulse energy (< 2 nJ) lasers, at 80 MHz, i.e. Ti:Sa and optical parametric oscillator (OPO, **i**). The Ti:Sa beam is separated by a beam splitter (BS) into two optical pathways, one being used to pump the OPO, the other being coupled into the microscope. Thus, the microscope covers a broad excitation range, i.e. 690-1080 nm (Ti:Sa) and 1050-1350 nm (OPO). Two types of optical parametric amplifiers (OPA), delivering high pulse energy radiation, are used as three-photon excitation sources. The Ytterbia OPA, a new OPA prototype, emits at a fixed wavelength (1650 nm, bandwidth 60 nm,) and variable repetition rate between 1 and 4 MHz and delivers up to 130 nJ pulse energy at the sample (**iii**). The state-of-the-art OPA is tunable between 1250 and 1700 nm, at 2 MHz, and delivers up to 40 nJ pulse energy at the sample (**ii**). Power control of laser pulses was performed using power attenuation (PA) units consisting of a half-wave plate and polarizing cube beam splitter. Due to the slightly positive group-velocity dispersion of our microscope, we built in a single-prism pulse compressor (PuCo) in the OPO beam path, and a ZnSe window in the Ytterbia OPA beam path. A flip mirror (FM) is used to accurately switch between OPA and OPO irradiation regimes. The Ti:Sa and OPO beam paths are overlapped before being coupled into the microscope, using a dichroic mirror (DM). For imaging, laser beams are scanned over the sample using a x/y galvanometric scanner. Fluorescence, higher-harmonics generation signals and excitation beams are separated using dichroic mirrors (DM), high-pass filters, and band-pass interference filters. Up to four photomultiplier tubes (PMT) are used for signal detection. The pulse trains of OPO and the two OPA systems are shown in **i-iii**. Box **iv** shows that imaging was performed in the medial region (crest) of the mouse tibia. Imaging requires the laser beam to surpass first the bone cortex and then the bone marrow, i.e. a soft lymphoid tissue, tissue compartments having distinct absorption and scattering properties.

**Table I:** Main parameters characterizing the microscope system: Maximum average power, maximum pulse energy, lateral (xy) and axial (z) dimensions of the point-spread function (PSF), i.e. dimensions of 100 nm fluorescent nanospheres embedded in agarose, measured using the Olympus lens XLPLN25XWMP2, NA 1.05.

| laser/wavelength | | rep. rate<br>(MHz) | max. av.<br>P/mW | max.<br>pulse<br>energy/<br>nJ | pulse<br>width / fs<br>-laser<br>output | pulse width<br>/ fs<br>-under the<br>lens | PSF xy /<br>nm | PSF z / nm | $\Delta\lambda$ /<br>nm |
| --- | --- | --- | --- | --- | --- | --- | --- | --- | --- |
| Ti:Sa | 930 nm | 80 | 720 | 9.0 | 140 | 247 | 368±27 | 1322±89 | 7 |
| OPO | 1100 nm | 80 | 230 | 2.875 | 200 | 166 | 407±42 | 1503±158 | 12 |
|  | 1330 nm | 80 | 66 | 0.825 | 200 | 166 | - | - | 16 |
| tunable<br>OPA | 1330 nm | 2 | 145 | 72.5 | 50 | 55 | 533±48 | 1537±167 | - |
|  | 1630 nm | 2 | 75 | 37.5 | 60 | 75 | 635±39 | 1700±175 | - |
| Ytterbia | 1650 nm | 1.01 | 144 | 142.6 | 65 | 72 | 587±109 | 1893±210 | 60 |
| OPA | 1650 nm | 2.06 | 286 | 138.8 | 65 | 71 | 691±31 | 1806±152 | 60 |
|  | 1650 nm | 3.09 | 376 | 130.7 | 65 | 73 | 684±72 | 1877±174 | 60 |
|  | 1650 nm | 3.98 | 568 | 142.7 | 65 | 75 | 669±55 | 1882±160 | 60 |

In contrast to the low-pulse-energy Ti:Sa and OPO systems (Fig. 1i), the OPA lasers deliver high-pulse-energy radiation to the sample (Fig. 1ii and iii), enabling efficient three-photon excitation and harmonics generation (*Suppl. Info.*). Thus, they are adequate for three-photon imaging in the mouse tibia (Fig. 1iv). The pulse energy values for selected wavelengths of the four lasers are provided in **Table I**.

### Three-photon excitation by high pulse energy 1650 nm radiation is required for high-resolution *in vivo* imaging through thick tibia cortex

Using our setup, we investigated the impact of excitation wavelength and pulse energy on the attenuation of radiation in bone and marrow tissue and, thus, on the imaging depth of *in vivo* multi-photon microscopy in long bones.

For this purpose, we performed *in vivo* 3D tibia imaging in adult Cdh5:tdTomato/ Histone:GFP reporter mice (hereafter Cdh5:tdTom) at 1650 nm (Ytterbia OPA), at 1330 nm (state-of-the-art OPA) and at 1100 nm (OPO). TdTomato fluorescence was detected using all excitation wavelengths, being induced at 1650 nm by non-resonant (simultaneous) three-photon excitation, at 1330 nm by resonant (sequential) three-photon excitation, i.e. two-photon excitation followed by one-photon excitation, and at 1100 nm by non-resonant two-photon excitation (Suppl. Fig. 2). GFP fluorescence was detected only at 1330 nm, upon non-resonant three-photon excitation (Suppl. Fig. 2). In Cdh5:tdTom reporter mice, tdTomato is expressed in the membrane, GFP in the nuclei of endothelial cells. 3D images were acquired over large volumes, i.e. up to 442×442 µm² (517×517 pixels), in the medial area of the tibia (Fig. 1iv) and up to 500 µm tissue depth, with step sizes of 2 or 4 µm.

As we expected stronger attenuation of radiation with increasing cortical bone thickness, we compared 3D images acquired in mice with intact, >100 µm thick tibia cortex (in average 136 µm, range 77 µm) and in mice with mechanically thinned cortex, <100 µm thick (in average 62 µm, range 28 µm) (Fig. 2a-d; Suppl. Fig. 3).

**Figure 2.**
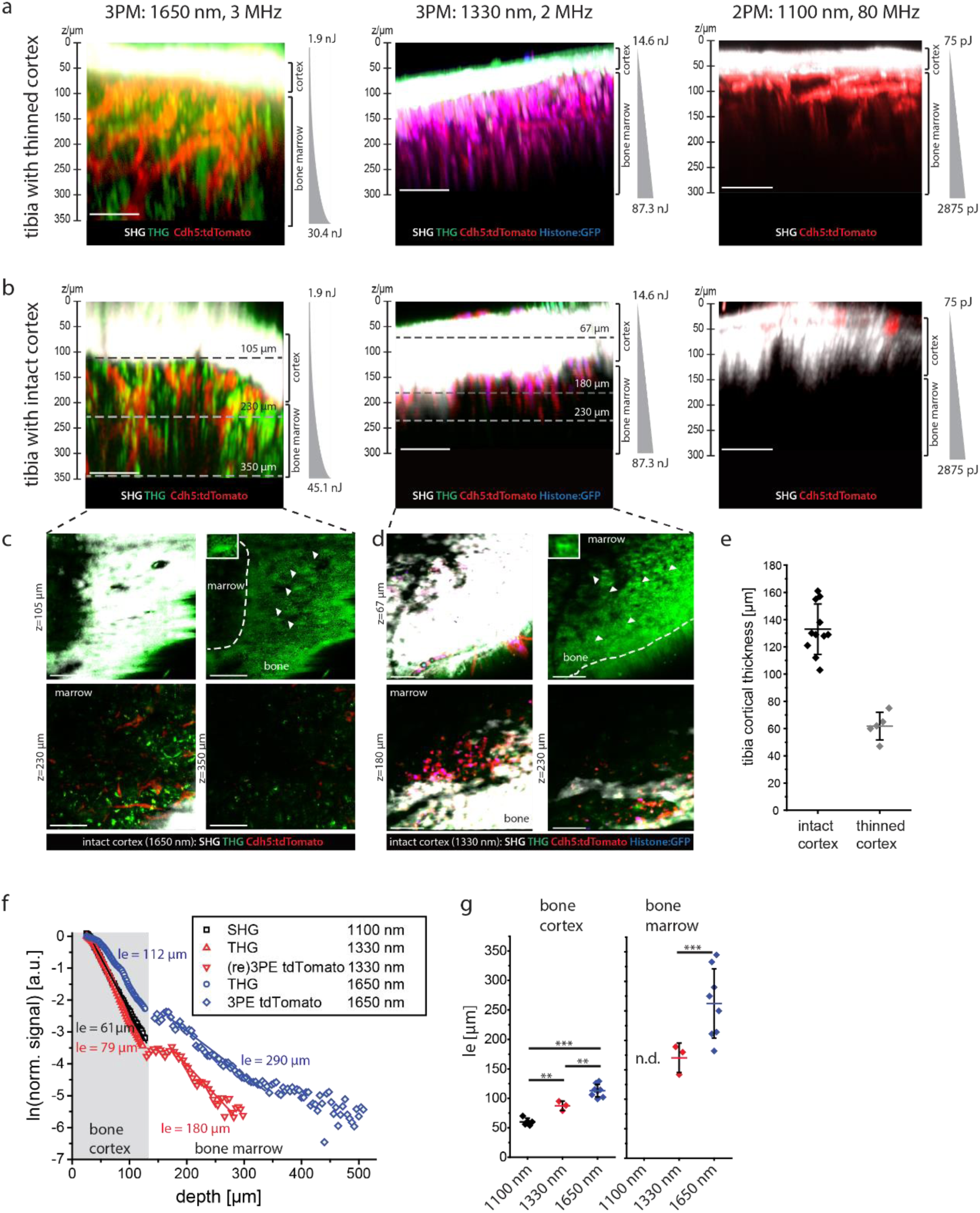
Minimizing attenuation of radiation by wavelength and pulse energy optimization for *in vivo* deep-marrow imaging in the intact mouse tibia, through >100 µm thick cortical bone. **a.** Axial xz 3D image projections acquired by three-(3PM) or two-photon microscopy (2PM) in tibia bones of Cdh5:tdTomato/ Histone:GFP (Cdh5:tdTom) mice, through mechanically thinned bone cortex, <100 µm thick. tdTomato and GFP in endothelial cells (vasculature) are shown in red and blue. Second harmonics generation (SHG) and third harmonics generation (THG) are shown in white and green. 3PM was performed either at 1650 nm, 3 MHz using the Ytterbia OPA (left) or at 1330 nm, 2 MHz using the tunable OPA (middle). 2PM was performed at 1100 nm, 80 MHz using the OPO (right). All excitation schemes enable bone marrow imaging, with imaging depths >350 µm at 1650 nm, ≈300 µm at 1330 nm and ≈200 µm at 1100 nm. **b**. Axial xz 3D image projections acquired by 3PM or 2PM in intact tibia bones of the same mouse strain, through >100 µm bone tissue. 3PM and 2PM were performed as indicated in **a**, showing that bone marrow imaging through thick bone is possible only by 3PM. Imaging depths >350 µm are achieved at 1650 nm, but only ≈ 230 µm (endosteal areas) at 1330 nm. 2PM at 1100 nm, 80 MHz enable only signal detection in bone cortex, not in the marrow. **a-b**. Indicated pulse energy and z-adaptation of power were chosen to prevent tissue damage. **c**. xy projections corresponding to the tissue layers indicated by dashed lines in **b**, left panel (3PM, 1650 nm). SHG and THG signals in bone cortex (105 µm depth) are shown in the upper panels. Arrowheads indicate THG signal in single lacunae (right), with an enlarged lacuna as inset. Blood vessels (tdTomato) and THG are shown in endosteal areas (230 µm depth) and in deep marrow (350 µm depth). **d**. xy projections corresponding the tissue layers indicated by dashed lines in **b**, middle panel (3PM, 1330 nm). Similar signals as in **c** are detected in bone cortex (67 µm depth) and in endosteal areas (180 µm, 230 µm depth), but not in deep marrow. **a-d**. Scale bar = 100 µm. **e**. Thickness of tibia cortex in the analyzed mice, either with mechanically thinned (n = 5 mice) or with intact cortex (n = 11 mice), determined relying on THG at 1330 nm and 1650 nm, and on SHG at 1100 nm. **f**. Effective attenuation length *l_e_* dependence on imaging depth z in the intact mouse tibia (>100 µm thick cortex). **g**. *l_e_* distribution in bone tissue and marrow (n = 5 mice at 1100 nm; n = 3 mice at 1330 nm; n = 8 mice at 1650 nm). n.d. – not detected. Statistical analysis was performed using two-way ANOVA with Bonferroni post-test or t-test, significance: * p > 0.05, ** p > 0.01, *** p > 0.001.

Imaging of both fluorescence and higher harmonics generation in the bone cortex and bone marrow of thinned tibia in Cdh5:tdTom mice was successfully performed using all excitation schemes (Fig. 2a). Through intact bone cortex thicker than 100 µm (Fig. 2b), imaging the tibia marrow was possible only upon excitation at 1650 nm (Fig. 2c) or at 1330 nm (Fig. 2d), with larger imaging depth at 1650 nm than at 1330 nm. Excitation at 1100 nm did not permit imaging through intact cortical bone thicker than 100 µm. The cortical thickness in each analyzed mouse (Fig. 2e) was determined based on the second harmonics generation (SHG) signal originating from collagen fibers in bone tissue, at 1650 nm, 1330 nm and 1100 nm. The results were validated using the third harmonics generation (THG) signal originating from lacunae harboring osteocytes, their connecting canaliculi and collagen fibers in bone cortex, at 1650 nm and 1330 nm.

Confirming these observations, imaging of marrow vasculature in the explanted tibia of over 100 weeks old Cdh5:tdTom female mice was possible at both 1100 and 1650 nm (Suppl. Fig. 4), as in old female mice the bone cortex is naturally thinner^50^.

To quantify the effect of excitation wavelength on the performance of deep-tissue tibia imaging, we assessed the effective attenuation length *l_e_* in bone cortex and bone marrow of Cdh5:tdTom mice with intact tibia at 1650 nm, 1330 and 1100 nm (Fig. 2f). According to Eq. (29)-(33) in *Suppl. Info.*, *l_e_* is a standardized measure for attenuation of radiation in tissue, defined as the tissue depth, at which the laser power decreases to 1/e of its value at the tissue surface (z = 0 µm).

At both 1650 nm and 1330 nm, the *l_e_* values in bone cortex are lower than in bone marrow (Fig. 2g), in agreement with a larger refractive index variation and a stronger scattering in calcified bone as compared to soft hematopoietic tissues^26,28^. The *l_e_* values determined for bone cortex are larger at 1650 nm excitation (113±11 µm, s.d.) when compared to 1330 nm (87±8 µm, s.d.) and to 1100 nm (60±6 µm, s.d.), in line with reduced scattering and lower signal attenuation at longer wavelengths^29^, as indicated in the previous section. The same holds true for the *l_e_* values measured in bone marrow, which were larger at 1650 nm (263±59 µm, s.d.) as compared to 1330 nm (170±25 µm, s.d.). In line with lower signal attenuation in lymphoid tissues compared to bone cortex, the *l_e_* values in bone marrow are larger than in bone tissue for both 1330 nm and 1650 nm. As imaging of tibia marrow through intact, thick bone cortex was not possible at 1100 nm, corresponding *l_e_* values cannot be determined. Notably, the interindividual variation of *l_e_* values in bone tissue is lower (value range 30 µm at 1650 nm) than in bone marrow (value range 163 µm at 1650 nm), consistent with a higher heterogeneity of tissue structure in the marrow as compared to bone cortex.

We expected that efficient three-photon excitation and third harmonics generation in deep-tissue tibia imaging require high pulse energy radiation. By comparing 3D images acquired in the intact tibia of a Cdh5:tdTom mouse *in vivo*, upon excitation at 1330 nm and average laser power 30 mW, at 80 MHz repletion rate and 2 MHz repetition rate, respectively, we showed that only excitation at high pulse energy (15 nJ at 2 MHz) but not at low pulse energy (0.38 nJ at 80 MHz) enables deep-marrow 3PM imaging in intact tibia (Suppl. Fig. 4).

Next to inducing attenuation of radiation, tissue scattering leads to the degradation of spatial resolution with increasing imaging depth, as compared to diffraction-limited resolution (dashed red lines in Fig. 3c, d). We assessed the depth-dependent degradation of spatial resolution in intact murine tibia (cortical thickness >100 µm) by analyzing xy- and z-profiles of THG signals upon excitation at 1650 nm (Fig. 3a, magenta line profiles in upper and lower close-up images, respectively; Video 1) and 1330 nm (Fig. 3b), respectively. Therefore, we approximated by Gaussian functions the xy- and z-profiles of the THG signal in canaliculi within bone cortex (black profiles), and in bright, granular structures present within cells in the bone marrow (red profiles, left graphs in both Fig. 3a and 3b). These results were confirmed by analyzing the first derivative of THG xy- and z-profiles, crossing edges of tissue structures both in bone cortex (black profiles) and marrow (red profiles), i.e. ∂THG/∂x and ∂THG/∂z, respectively (right graphs in Fig. 3a and 3b). The depth-dependent degradation of spatial resolution was more accentuated at 1330 nm than at 1650 nm excitation, in both bone cortex and bone marrow (Fig. 3c, d). In the endosteal region, we measured 0.6±0.1 µm lateral (upper panel Fig. 3c) and 2.2±0.2 µm axial resolution (lower panel Fig. 3c) at 1650 nm, and 0.9±0.2 µm (upper panel Fig. 3d) and 2.8±0.2 µm (lower panel Fig. 3d) at 1330 nm. In the bone marrow, the lateral (upper panels Fig. 3c and 3d) and axial resolution (lower panels Fig. 3c and 3d) amounted to 0.9±0.1 µm and 2.6±0.2 µm at 1650 nm (in 300 µm tissue depth), and to 1.4±0.2 µm and 3.8±0.5 µm at 1330 nm (in 270 µm tissue depth).

**Figure 3.**
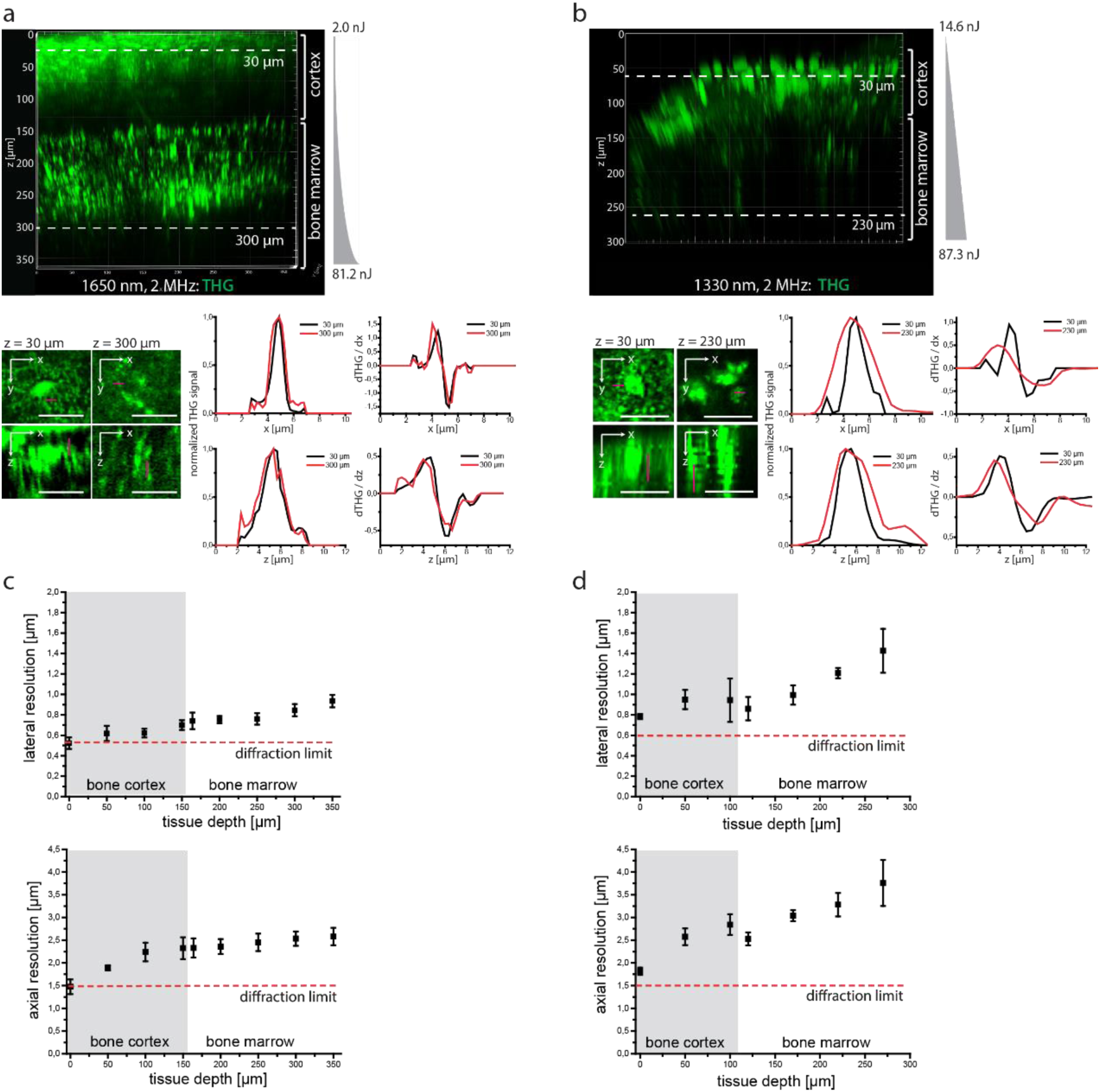
Three-photon imaging at 1650 nm enables sub-cellular resolution throughout cortical bone and bone marrow in intact tibia. **a**. 3D reconstruction (442×442×362 µm³, 1036×1036×362 voxel) of THG in the intact tibia, at 1650 nm, 2 MHz (upper panel). Representative xy and xz projections of THG in 30 µm depth in bone cortex and in 300 µm depth in bone marrow (bottom image array), tissue layers indicated in the upper panel. Scale bar = 30 µm. Representative intensity profiles of THG signal (left, corresponding to the magenta lines in the xy and xz projections) and their first derivatives (right), measured in 30 µm (black profiles) and 300 µm tissue depth (red profiles). **b**. 3D reconstruction (442×442×300 µm³, 517×517×150 voxel) of THG in the intact tibia, at 1330 nm, 2 MHz (upper panel). Representative xy and xz projections of THG in 30 µm depth in bone cortex, and in 230 µm depth in the endosteal area, as indicated in the upper panel. Scale bar = 30 µm. Similar to a, THG intensity profiles (left, corresponding to the magenta lines in the xy and xz projections) and their first derivatives (right) are shown in 30 µm (black profiles) and 230 µm depth (red profiles). **a-b**. Pulse energy and z-adaptation of power are indicated. **c**. Depth dependence of lateral and axial resolution determined in the intact tibia upon excitation at 1650 nm, based on Gaussian approximation of THG intensity profiles and their first derivatives. For each tissue depth, at least 5 x- and 5 z-profiles were averaged (error bars show s.d.). **d**. Depth dependence of lateral and axial resolution determined in the same manner as described for **c**, at 1330 nm. **c-d**. Axial and lateral resolution deteriorate with increasing imaging depth at both 1650 nm and 1330 nm excitation, with less degradation at 1650 nm. Thus, 3PM at 1650 nm preserves subcellular resolution in the marrow cavity, with lateral resolution values better than 1 µm and axial resolution values of ≈2.5 µm, in 300 µm depth. The diffraction limit of microscope was calculated based on the paraxial approximation, confirmed by 3PM of fluorescent nanospheres, at both 1650 nm and 1330 nm (**Table I**), and displayed as dashed red lines.

Thus, we concluded that high pulse energy laser radiation of 1650 nm wavelength is required for *in vivo* deep-tissue tibia imaging through intact bone cortex thicker than 100 µm.

### Repetition rate optimization of 1650 nm radiation facilitates fast *in vivo* deep-marrow imaging in intact tibia

The average number of laser pulses per pixel rises with elevated laser repetition rate (graphs in Fig. 4a), leading to a linear increase of detected signal (fluorescence, SHG or THG), if pulse energy and pixel dwell time are kept constant. Conversely, for the same signal quality, higher repetition rates allow faster time-lapse imaging.

**Figure 4.**
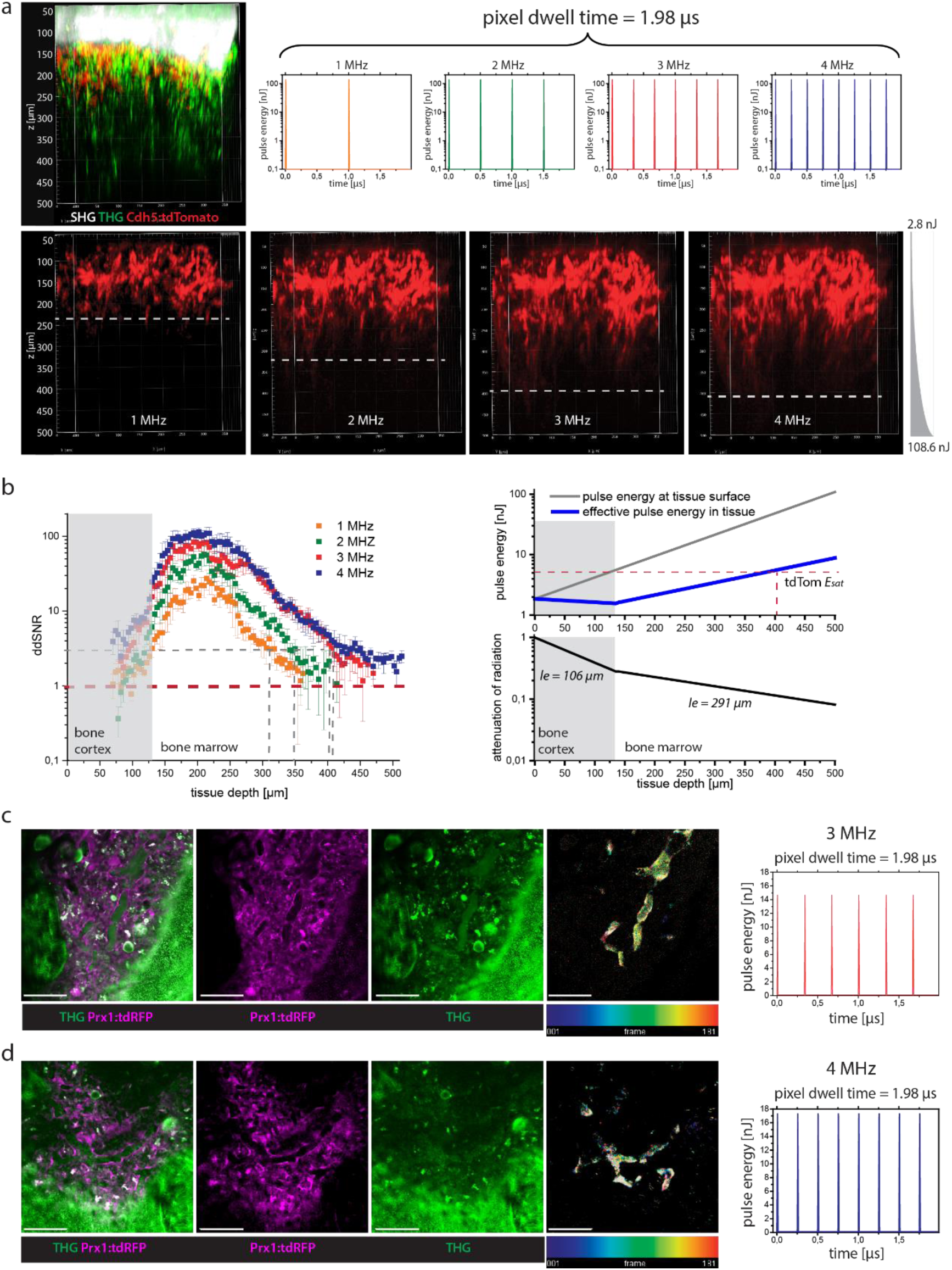
Laser repetition rate optimization for fast image acquisition in deep tissue layers of intact tibia by three-photon excitation at 1650 nm *in vivo*. **a.** 3D reconstructions (400×400×500 µm³, 518×518×125 voxel) of tdTomato (red), SHG (white) and THG (green) in the intact tibia of a Cdh5:tdTom mouse at 1650 nm, with 1, 2, 3 and 4 MHz. Deeper imaging was achieved at higher repetition rates, as indicated by the dashed lines. The pulse trains at each repetition rate for 1.98 µs pixel dwell time are shown in the graphs. **b**. *Left panel*: Depth dependent SNR determined for tdTomato fluorescence at 1 MHz (orange), 2 MHz (green), 3 MHz (red) and 4 MHz (blue). The absolute detection limit is given by SNR = 1 (red line). SNR = 3 is needed for reliable 3D object segmentation (grey line), being reached in 300 µm depth at 1 MHz, 340 µm at 2 MHz and ≈400 µm at both 3 and 4 MHz. *Right panel*: Depth dependent applied pulse energy at the tibia surface (grey line) and effective pulse energy in tissue (blue line), upper graph. The effective pulse energy is the product of the pulse energy at the tibia surface and of the normalized attenuation of radiation in tissue (bottom graph). **c**. First xy image (400×400 µm², 518×518 pixel) of a time-lapse 2D stack acquired by *in vivo* 3PM in the tibia of a Prx1:tdRFP mouse at 1650 nm, 3 MHz (pulse energy 16 nJ, in 126 µm tissue depth, pulse train shown in graph). **d**. First xy image of a similar time-lapse 2D stack as in **c** acquired at 4 MHz (pulse energy 14 nJ, in 120 µm depth, pulse train shown in graph). **c-d**. tdRFP fluorescence (stroma compartment) is shown in magenta, THG in green. Among other tissue components, erythrocytes show THG signal, enabling to visualize blood flow in a label-free manner. Videos were acquired over 3 minutes, every second. To generate the time color-coded image (right images), we calculated the difference between every two consecutive THG images, color-coded the resulting images according to the acquisition time-point and summed them up. In this way, only regions with changing structures, such as blood flow, are highlighted. 3PM at both 3 and 4 MHz enables blood flow visualisation over large fields of view in the tibia marrow. As at the same pulse energy, the average laser power at 3 MHz is lower than at 4 MHz, imaging at 3 MHz is less prone to induce tissue photodamage. Scale bar = 100 µm.

We took advantage of the particular properties of the novel OPA design, i.e. variable repetition rate up to 4 MHz and high pulse energy, similar at all repetition rates (**Table I**), to investigate the effect of higher repetition rates and, thus, of higher signal strength, on imaging depth and image acquisition speed in the intact tibia *in vivo*.

First, we performed *in vivo* 3D imaging of intact tibia in a Cdh5:tdTom mouse, with 130 µm thick bone cortex, at 1650 nm and variable repetition rates, keeping pixel dwell time at a value of 1.98 µs, with 2x frame averaging (Fig. 4a; Video 2). For all four repetition rates, we used the same exponential z-adaptation of pulse energy (2.76 nJ at z = 50 µm below the cortex surface, and 108.64 nJ at z = 500 µm tissue depth). We detected tdTomato fluorescence of endothelia in deeper layers of the tibia marrow, at 3 and 4 MHz repetition rate, as compared to 1 and 2 MHz, consistent with an increased number of laser pulses arriving at the sample within the same pixel dwell time (Fig. 4a).

To quantify this observation, we determined the depth-dependent signal-to-noise ratio (ddSNR) of the tdTomato fluorescence signal for all repetition rates (Fig. 4b, left panel). SNR (Eq. (36), *Suppl. Info.*) is defined as the ratio between the difference of measured fluorescence signal and mean background value and the width of the background count distribution, i.e. the noise. When the SNR value equals 1, the signal cannot be distinguished from noise and therefore the maximum imaging depth is reached. We found that the ddSNR of tdTomato fluorescence decreased in the same manner for all laser repetition rates. As the SNR value scales up linearly with the repetition rate, for higher repetition rates the ddSNR reached the value of 1 in deeper marrow layers and, thus, the maximum imaging depth increased (SNR = 1 is marked as dashed red line in Fig. 4b). From our experience, reliable object segmentation, necessary for cell tracking, characterization and classification, requires SNR values of at least 3 (dashed grey lines in Fig. 4b), in agreement with previous reports^39^. An SNR value of 3 was reached in ≈400 µm depth both at 3 MHz and 4 MHz repetition rate, deeper than in ≈300 µm at 1 MHz and in ≈350 µm at 2 MHz.

The imaging depth at 1 and at 2 MHz repetition rate can be increased either by increasing the pixel dwell time but impairing the image acquisition speed. Another strategy is to use a steeper z-adaptation of pulse energy at 1 or 2 MHz as compared to 3 and 4 MHz, at the same incremental average power, i.e. the same exposure to thermal damage. However, as the applied increase of pulse energy with depth (Fig. 4b, right panel) was chosen in such a way that the saturation energy *E_sat_* of tdTomato (4.03 nJ, quantification described in *Suppl. Info.*) is reached in 400 µm depth, a steeper increase of pulse energy does not lead to fluorescence signal increase, but to highly non-linear photobleaching and, possibly, to non-linear phototoxicity. According to the z-adaption of power used in our experiment, the pulse energy applied to image marrow layers in 400 µm depth was 39.8 nJ (average laser power 119 mW at 3 MHz and 158 mW at 4 MHz).

Whilst excitation at 3 and 4 MHz repetition rates permitted similar maximum imaging depths in the intact tibia, the average power at 4 MHz is higher at the same pulse energy, which may lead to thermal and photodamage of tissue. Hence, we expected a repetition rate of 3 MHz to be more advantageous for *in vivo* time-lapse tibia imaging experiments, if it allows the visualization of fast biological processes, such as blood flow.

We performed *in vivo* time-lapse imaging of 400×400 µm² areas (518×518 pixels) in the tibia of Prx1:tdRFP fate mapping mice at 1650 nm, at both 3 and 4 MHz, over 3 minutes, every second, i.e. with 1 Hz acquisition rate and pixel dwell time 1.98 µs (Fig. 4c,d; Video 3,4). In Prx1:tdRFP fate mapping mice, mesenchymal stromal cells and their progeny express tdRFP. The pulse energy and average power values used for image acquisition together with the location of the imaged tissue layer in the tibia are provided in **Table II**. TdRFP fluorescence appeared throughout the marrow tissue, spearing vessel lumina, whereas THG signal was detected in both parenchyma and the vasculature (Fig. 4c,d). Inside blood vessels, THG signal stemming from erythrocytes allowed us to visualize blood flow at distinct flow rates in different vessel types (Video 3,4), in line with previous reports^51^. By reducing the size of the field of view (200×200 µm², 257×257 pixels, pixel dwell time 4 µs), we achieved even faster imaging at 2 Hz acquisition rate, i.e. a frame every 0.5 s, at both 3 and 4 MHz (last part in Video 3,4).

**Table II:** Parameter overview for *in vivo* time-lapse 2D imaging experiments in the tibia of Prx1:tdRFP mice (n = 2).

| exc. scheme | time step [s] | acq. time [min] | cortex thickness [μm] | $l_e$ [μm] | marrow thickness [μm] | $l_e$ [μm] | pulse energy at surface [nJ] | eff. pulse energy [nJ] | power at surface [mW] | pixel dwell time [μs] | |
| --- | --- | --- | --- | --- | --- | --- | --- | --- | --- | --- | --- |
| 1650 nm, 3 MHz | 1 | 3 | 96 | 89 | 30 | 259 | 16 | 4.9 | 50 | 1.98 | Fig. 4; Video 3 |
| 1650 nm, 4 MHz | 1 | 3 | 80 | 100 | 40 | 291 | 14 | 5.5 | 57 | 1.98 | Fig. 4; Video 4 |

Concluding, we found similar performance at both 3 and 4 MHz repetition rates for fast *in vivo* marrow imaging. In the following experiments we decided to employ excitation at 1650 nm and 3 MHz repetition rate, not exceeding 120 mW average power (39.8 nJ pulse energy), conditions we found necessary for imaging down to 400 µm depth in the marrow cavity of intact tibia. Next, we investigated whether these experimental conditions lead to thermal or photodamage of the tissue or resulted in strong fluorophore photobleaching.

### Negligible tissue photodamage and tdRFP photobleaching during *in vivo* time-lapse tibia imaging at 1650 nm, 3MHz

Tissue photodamage induced by pJ to nJ pulse energy radiation, in the near-infrared and infrared range, is a complex process thought to originate from two main components: the bulk heating effect of infrared radiation throughout the exposed tissue, which scales linearly with the average laser power^52^, and non-linear photodamage effects at the focal plane, induced by the high density of pulse energy^51^. Due to the attenuation of radiation, the sample surface is most endangered by overheating, as it is exposed to a higher laser power than the underlying tissue.

To rule out overheating of the bone tissue during *in vivo* time-lapse deep-marrow imaging of intact tibia in mice, we set the average laser power of 1650 nm radiation, at 3 MHz repetition rate to 120 mW and performed repeated imaging of 400×400×30 µm³ volumes in the tibia marrow over 2 hours, every 30 s, in 300-350 µm tissue depths. We found no changes of either THG or SHG signals in the cortical bone layers above the imaged site in the tibia marrow, before and after *in vivo* time-lapse imaging (Suppl. Fig. 5; *Suppl. Info.*). Furthermore, no signs of tissue damage were detected at the bone surface of 3PM imaged tibias using Nanofocus-CT, scanning electron microscopy and Movat’s pentachrome staining, compared to similar surface areas of non-irradiated contralateral bones (Suppl. Fig. 5; *Suppl. Info.*). Thus, we concluded that 120 mW average laser power at 1650 nm and 3 MHz radiation does not affect thermally the bone tissue and is biocompatible for time-lapse *in vivo* imaging of intact tibia.

Next, using immunofluorescence analysis, we investigated possible damage at the imaged tissue site (300-350 µm total tissue depth) in the marrow of intact tibia bones induced by time-lapse *in vivo* imaging in CD19:tdRFP fate mapping mice, at 1650 nm, 3MHz (120 mW average power, 39.8 nJ pulse energy). In CD19:tdRFP fate mapping mice, tdRFP is expressed in B lineage cells. Time-lapse imaging was performed over 2 hours, every 30 s. Thereafter the imaged bones were explanted and prepared for immunofluorescence analysis, using various antibodies indicative of photodamage (Fig. 5a). The non-irradiated contralateral tibia bones were used as controls.

**Figure 5.**
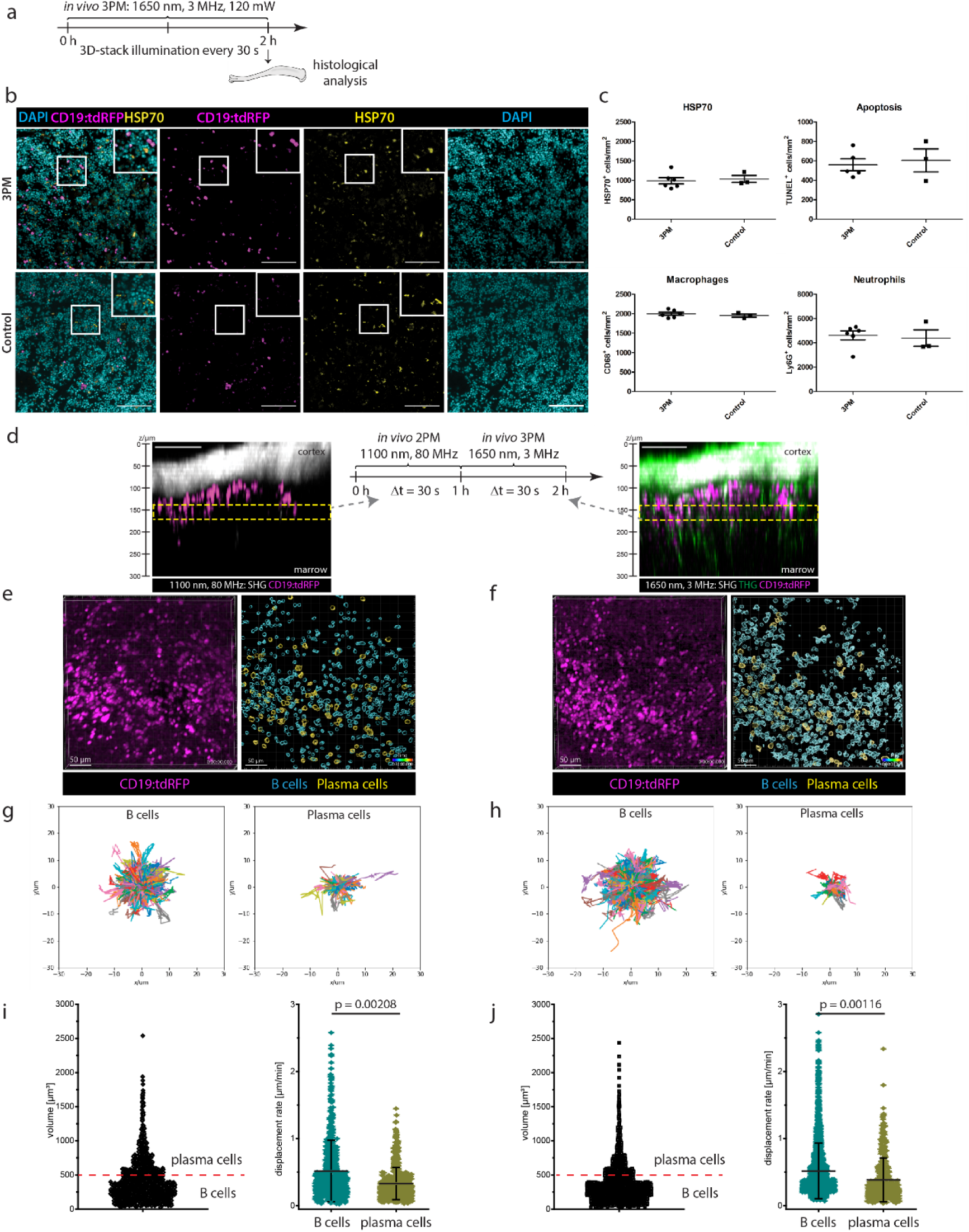
Multimodal tissue analysis reveals no signs of photodamage by *in vivo* time-lapse three-photon imaging at 1650 nm in mouse tibia. **a.** Experimental design for immunofluorescence histological analysis of marrow tissue after intravital 3PM at 1650 nm, 3 MHz in the tibia of CD19:tdRFP mice. 3D-imaging was performed over 2 hours, every 30 s, at 40 nJ (120 mW). **b**. Representative immunofluorescence overlays of B lineage cells (tdRFP, magenta), heat-shock protein (HSP70, yellow) and nuclear staining (DAPI, cyan) in tibia marrow tissue irradiated by 3PM and not irradiated (control). Scale bar = 100 µm. **c**. Photodamage quantification by immunofluorescence analysis: frequencies of HSP70^+^ cells, apoptotic cells (TUNEL^+^), macrophages (CD68^+^) and neutrophil granulocytes (Lys6G^+^) in 3PM irradiated marrow tissue (n = 6 mice) are similar to controls (n = 3 mice). This indicates no signs of tissue photodamage by 3PM. **d**. Experimental design of *in vivo* imaging to compare the effect of 3PM to state-of-the-art 2PM on the motility of marrow B lineage cells. The same marrow tissue site in the tibia of CD19:tdRFP mice, with thinned cortex, was imaged by time-lapse 2PM at 1100 nm, 80 MHz, followed by time-lapse 3PM at 1650 nm, 3 MHz. Yellow rectangles indicate the repeatedly imaged volume in z projections acquired by 2PM (left) and 3PM (right). B lineage cells (tdRFP) are shown in magenta, SHG in white, THG in green. Scale bar = 100 µm. **e-f**. Representative 3D images (400×400×30 µm³, 518×518×11 voxel) of marrow B lineage cells (tdRFP) acquired by time-lapse 2PM (**e**) and 3PM (**f**) (left images). Corresponding results of tdRFP^+^ cell segmentation (right images). tdRFP^+^ cells with a cellular volume between 65 and 500 µm³ are defined as B cells (cyan), those with a volume between 500 and 4189 µm³ as plasma cells (yellow). Scale bar = 50 µm. **g-h**. Rose plots representing the cell tracks of B cells (left; n = 1121 cells for 2PM and n = 2006 cells for 3PM, in the same mouse) and plasma cells (right; n = 56 cells for 2PM and n = 136 cells for 3PM) over 30 min (2PM in **g**, 3PM in **h**). **i-j**. Cell volume distribution of segmented tdRFP^+^ cells from the time-lapse data (left). Volume threshold of 500 µm³ (cell diameter 10 µm) is indicated by the red line. Mean displacement rate distributions of B cells and plasma cells, respectively (right). 2PM data are shown in **i**, 3PM data in **j**. **d-j**. As the cell motility behavior of both marrow B cells and plasma cells is similar when analyzed by 2PM and by 3PM, we conclude that 3PM at 1650 nm, 3 MHz is reliable to assess cell dynamics *in vivo*. Statistical analysis was performed using t-test, p values indicated.

No increase in the number of cells expressing heat shock protein 70 (HSP70)^31^, a marker of photodamage, was observed at marrow tissue sites irradiated at 1650 nm compared to non-irradiated tibia marrow (Fig. 5b,c). Additionally, there was no evidence of increased apoptosis analyzed by TUNEL staining, or in the number of neutrophils (Ly6G), or macrophages (CD68), which are known to respond in a rapid manner to (laser) damage, by migrating into the respective tissue regions^53^ (Fig. 5c, Suppl. Fig. 6). In contrast, clear signs of tissue photodamage were detected by immunofluorescence analysis of spleens repeatedly exposed *in vivo* to excessive irradiation at 850 nm, 80 MHz, average power >300 mW (*Suppl. Info.*; Suppl. Fig. 6). A similar control experiment was not possible in intact tibia, as 850 nm radiation do not penetrate through thick bone cortex.

To confirm that repeated exposure to 1650 nm, 3 MHz laser radiation does not induce tissue photodamage, we performed in the thinned tibia of CD19:tdRFP mice first *in vivo* time-lapse 2PM at 1100 nm, 80 MHz followed by *in vivo* time-lapse 3PM at 1650 nm, 3 MHz at the same marrow site (Fig. 5d). For both excitation schemes, time-lapse imaging of marrow tdRFP^+^ B lineage cells was performed over 60 minutes, every 30 s (Video 5), and their motility patterns were analyzed. To that end, noise was removed in the acquired 3D images using a trained Noise2Void algorithm^54^, tdRFP^+^ cells were segmented (Fig. 5e,f) and tracked over time (Fig. 5g,h), as detailed in *Methods*. Pulse energy, average power and location of the imaged site in the medial region of tibia are summarized in **Table III** for both excitation schemes. The tdRFP^+^ B lineage cell population in the bone marrow comprises both B cells and plasma cells, as tdRFP expression is preserved also after down-regulation of CD19 expression at later stages of B lymphocyte differentiation. We distinguished between B cells and plasma cells among the marrow tdRFP^+^ cells relying on their cell volume. We assumed a cell diameter of 5 to 10 µm for B cells (V < 500 µm³) and of 10 to 20 µm for plasma cells (V > 500 µm³) (Fig. 5i,j), as previously reported^8,24^. The motility characteristics of marrow tdRFP^+^ cells determined by 2PM and by 3PM were similar, with a mean velocity of 3.64±0.76 µm/min (s.d.) and of 3.77±0.79 µm/min (s.d.), respectively.

**Table III:**
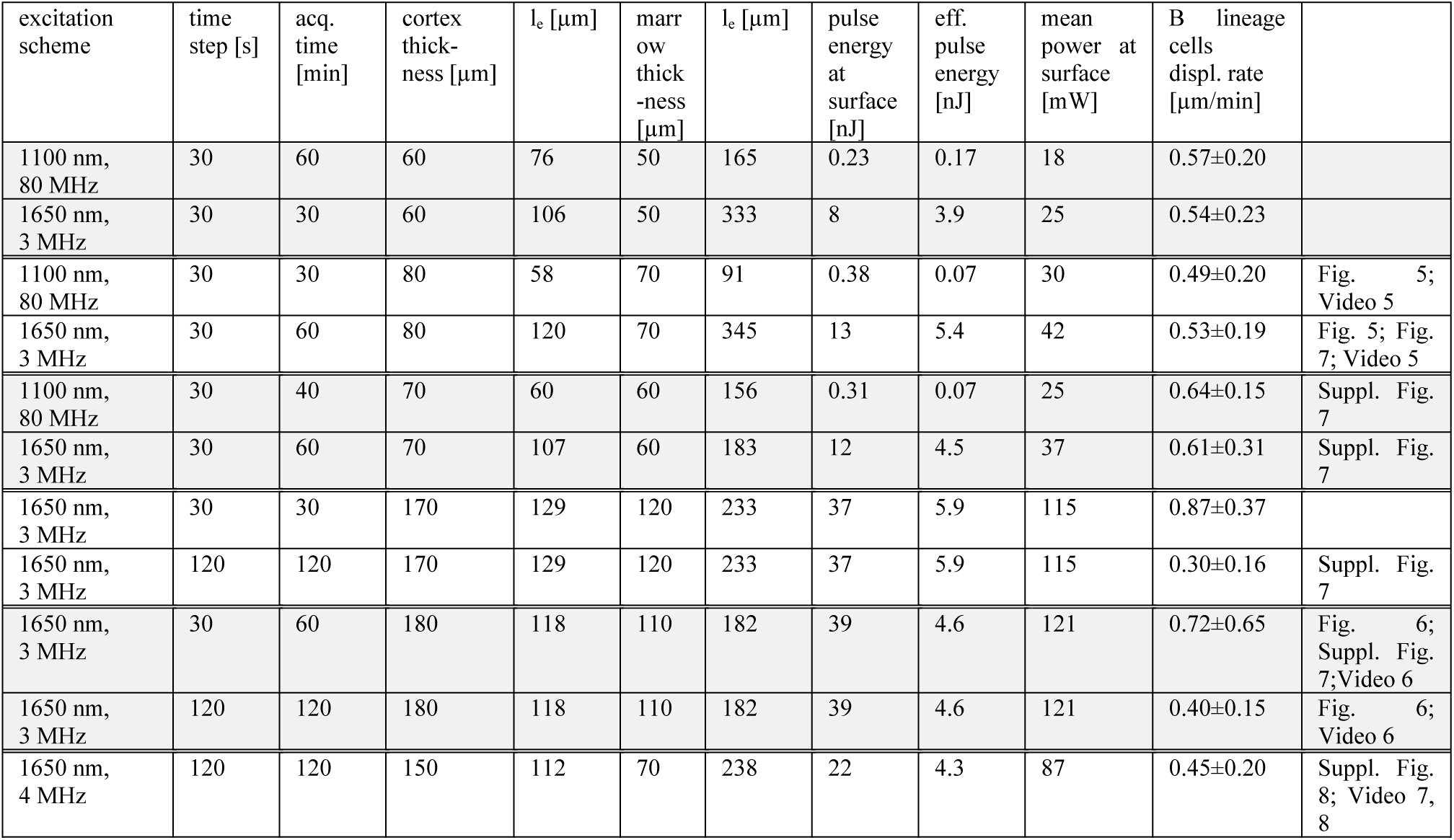
Parameter overview for *in vivo* time-lapse 3D imaging experiments in the tibia of CD19:tdRFP mice (n = 6).

In previous reports, we and others demonstrated that B cells are more motile in the bone marrow than plasma cells. For both excitation schemes (1100 nm and 1650 nm excitation), we confirmed this finding, as highlighted by B cell and plasma cell tracks acquired over 30 minutes (rose plots in Fig. 5g,h). We found mean displacement rates of 0.52±0.23 µm/min (s.d.) for B cells and 0.36±0.12 µm/min (s.d.) for plasma cells by 2PM, and of 0.55±0.20 µm/min (s.d.) for B cells and 0.38±0.16 µm/min (s.d.) for plasma cells by 3PM (Fig. 5i,j). We confirmed these findings in two replicate experiments (**Table III**). The mean displacement rate values are in agreement with previous reports^8,24,55^. Lower B lineage cell displacement rates were measured if time-lapse imaging was performed every 120 s as compared to every 30 s, as at lower sampling rates fast cells (typically B cells, not plasma cells) are missed by the tracking algorithm.

Next, we analyzed the tdRFP photobleaching caused by *in vivo* 3PM at 1650 nm, 3MHz compared to 2PM at 1100 nm, 80 MHz (Suppl. Fig. 7; *Suppl. Info.*). We found similar photobleaching rates of tdRFP fluorescence for both excitation schemes (*k_photobl_* = 2.4·10^-3^ min^-1^ at 1100 nm, 25 mW and *k_photobl_* = 1.3·10^-3^ min^-1^ at 1650 nm, 37 mW, Suppl. Fig. 7). At 1650 nm, 3 MHz, the tdRFP photobleaching rate increased with higher image acquisition rate but remained at a low level (Suppl. Fig. 7; *Suppl. Info.*). Owing to undetectable tissue photodamage and negligible fluorophore photobleaching, we concluded that the proposed three-photon microscopy method which uses 1650 nm, 3 MHz laser radiation at up to 40 nJ pulse energy is suitable to study cellular dynamics in the deep marrow of intact long bones *in vivo*, over large imaging volumes. This ability is particularly relevant when analyzing rare cell populations *in vivo*, such as the marrow plasma cells.

### Cell motility and label-free THG signal describe heterogeneity among marrow plasma cells *in vivo*

To analyze the dynamics of marrow plasma cells *in vivo*, we performed time-lapse deep-marrow 3PM (400×400×30 µm³, 518×518×11 voxels) in the intact tibia of CD19:tdRFP mice over a total of 3 hours, at 1650 nm, 3 MHz excitation (Fig. 6a,b; average power and pulse energy in **Table III**). Within the first hour, we acquired a 3D image every 30 s, and in the following 2 hours every 120 s (Fig. 6a,c; Video 6). Again, we found B cells to show larger displacement than plasma cells over a time period of 2 hours (rose plots in Fig. 6d). Notably, the mean displacement rate distribution of marrow plasma cells, i.e. tdRFP^+^ B lineage cells with a cell volume between 500 and 4189 µm³, could be best approximated by a bi-modal Gaussian function (Fig. 6e). This indicates two plasma cell subsets with distinct migratory behavior, 37.8% of the marrow plasma cells having a higher mean displacement rate of 0.96±0.38 µm/min than 62.2% with a mean displacement rate of 0.21±0.12 µm/min (n = 176 tracked plasma cells within an imaging volume). We could confirm our results in three independent experiments (Fig. 6f; Suppl. Fig. 8; Video 7). Our findings are in line with a report by Benet et al.^5^, which showed both highly-motile and non-migratory plasma cells by means of 2PM in the thinned tibia of Blimp1 reporter mice.

**Figure 6.**
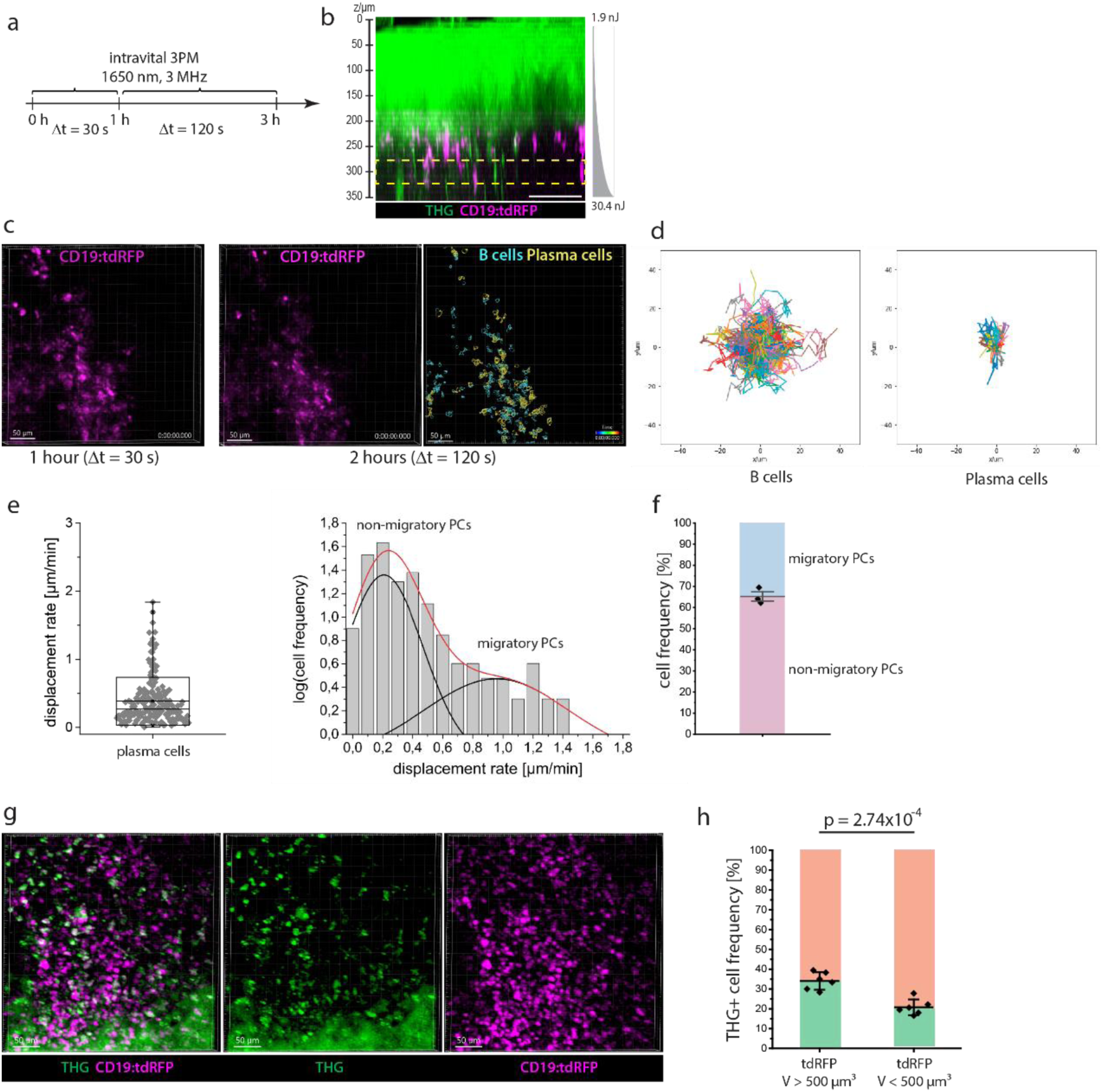
Heterogeneity within plasma cell population in the tibia marrow is defined by both their cellular migration patterns and THG signal. **a**. Experimental design of *in vivo* time-lapse 3PM at 1650 nm, 3 MHz in the intact tibia of CD19:tdRFP mice to study the migratory behavior of marrow plasma cells. A first acquisition period of 1 hour, every 30 s is followed by acquisition over 2 hours, every 120 s. **b**. z projection of intact tibia acquired by 3PM at 1650 nm, 3 MHz (exponential pulse-energy z-adaptation: 1.9 to 30.4 nJ). THG is shown in green, B lineage cells (tdRFP) in magenta. The yellow rectangle indicates the repeatedly imaged volume (≈300 µm depth). Scale bar = 100 µm. **c**. Representative 3D images of B lineage cells (400×400×30 µm³, 518×518×11 voxel) acquired during the first (left) and second imaging period (middle). Result of B lineage cell segmentation based on the data acquired during the second imaging period (2 h, right). B cells (cyan) and plasma cells (yellow) were defined by volume. Scale bar = 50 µm. **d**. Rose plots representing cell tracks of B cells (left; n = 765 cells) and plasma cells (right; n = 64 cells). **e**. Mean displacement rate distribution of marrow plasma cells assessed from the cell tracks measured over the entire 3 hours imaging period in a single mouse (left). Logarithmic representation of cell frequency histogram with respect to the mean displacement rate (right), fitted by a double Gauss-peak distribution, indicating two distinctly motile plasma cell subsets in the tibia marrow. The black lines represent the Gaussian fitting functions, the red line represents their sum. **f**. Relative cell frequencies of migratory and non-migratory marrow plasma cells in n = 3 CD19:tdRFP mice. **g**. Merged (left) and single channel (middle, right) 3D images (400×400×100 µm³, 518×518×51 voxel) of tdRFP fluorescence (magenta) in B lineage cells and THG (green) in the intact tibia of a CD19:tdRFP mouse, by *in vivo* 3PM at 1650 nm, 3 MHz, 21 nJ pulse energy, showing heterogeneous THG signal distribution among B lineage cells. Scale bar = 50 µm. **h**. Percentage of THG^+^ cell numbers within the B cell and plasma cell population, respectively (n = 6 mice). Whereas THG signal is enriched in marrow plasma cells as compared to B cells, only 1/3 of the detected plasma cells display the signal. Statistical analysis was performed using t-test, p value indicated.

By additionally analyzing the label-free THG signal in the acquired *in vivo* time-lapse 3D images within the tibia marrow of CD19:tdRFP mice (Fig. 6g), we found enriched THG signal in plasma cells as compared to B cells (Fig. 6h). For quantification, we segmented tdRFP^+^ cells and distinguished between B cells and plasma cells based on their volume (65µm³<V<500 µm³ and 500µm³<V<4189 µm³, respectively), as previously described^8^ and we analyzed the presence of a strong THG signal, defined as highest signal appearing in 1% of voxels, within the segmented cells. Despite an enrichment of THG signal in marrow plasma cells, only a minority (34.4±3.3%) of these cells contained THG signal (Fig. 6h).

Next, we aimed to answer whether the heterogeneous distribution of THG signal in marrow plasma cells is related to the biological function of these cells and if it correlates with their motility patterns.

### THG^hi^ signal in marrow plasma cells links antibody production capacity to cellular motility

In line with previous reports^40,44^, we observed a heterogeneous intensity distribution of THG both in tibia cortex and marrow by *in vivo* 3PM upon 1650 nm excitation. By analyzing the THG intensity both in bone cortex and bone marrow, we were able to distinguish between THG^lo^ and THG^hi^ signals (Fig. 7a, left panel). In the tibia marrow, the THG^lo^ signal mainly originated from cell membranes (lipid bilayers), whereas THG^hi^ stemmed from granules inside cells, presumably the lipid bilayers of cell organelles (Fig. 7a(i)). This observation is in line with a previous study showing a higher granularity associated with strong THG signal in leukocytes^44^. The THG^hi^ signal in bone cortex was associated mainly with lacunae, harboring osteocytes^56^ (Fig. 7a(ii)). Thin canaliculi connecting the lacunae showed a THG^lo^ signal. In the tibia marrow, we were able to use both THG^hi^ and THG^lo^ signals to segment single cells (cyan outlines for THG^lo^ cells, yellow outlines for THG^hi^ cells in Fig. 7b). Time-lapse 3D imaging of these THG signals enabled monitoring of whole-tissue dynamics at cellular resolution, in a label-free manner (Video 8) and, thus, represents a promising strategy to assess tissue strain as well as shear stress in bone and bone marrow.

**Figure 7.**
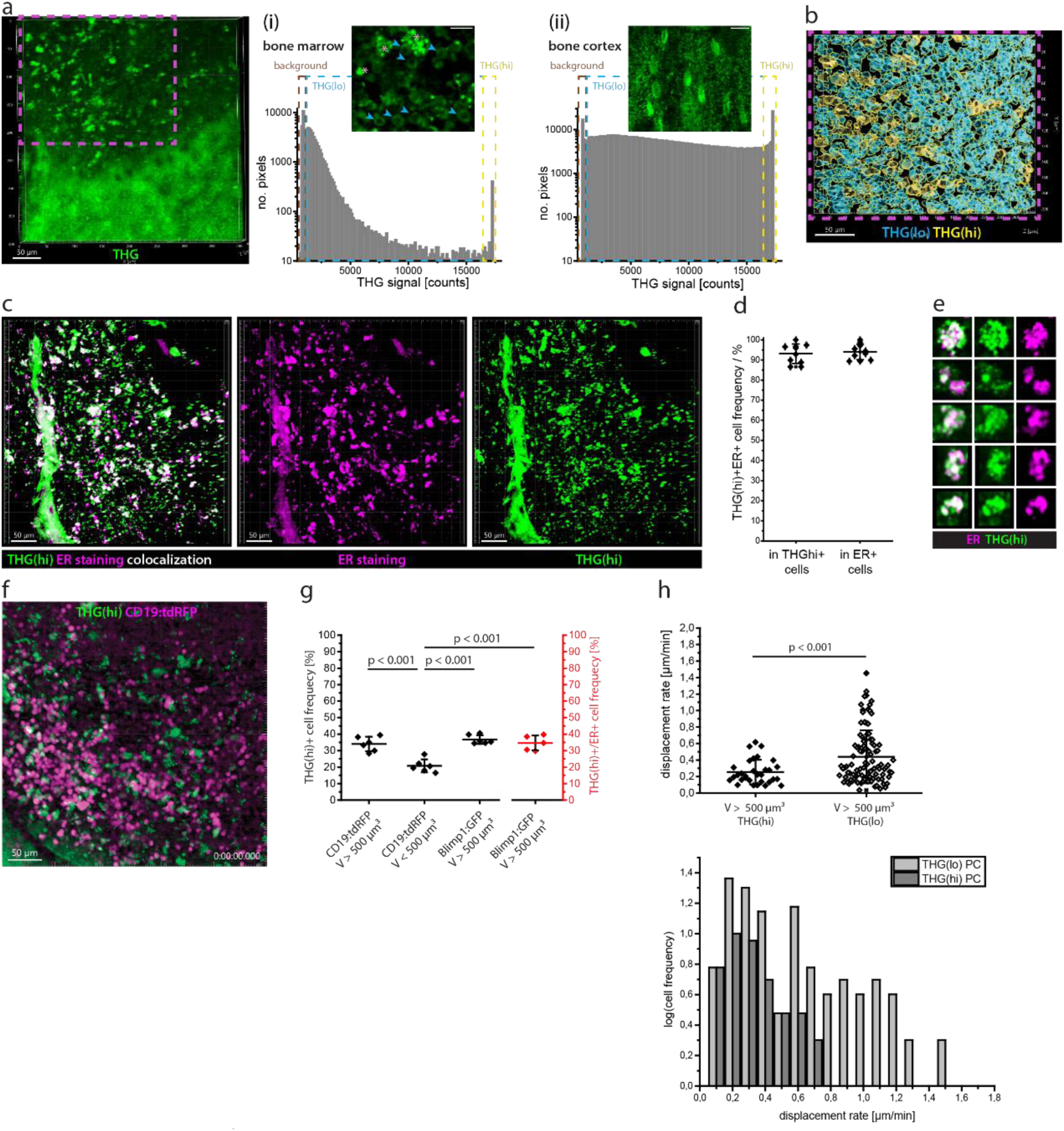
THG^hi^ signal defines two marrow plasma cell subsets with distinct functional capacity and migratory behavior. **a**. Representative 3D image (400×400×100 µm³, 518×518×11 voxel) of THG signal (green) in tibia cortex and marrow acquired by *in vivo* 3PM at 1650 nm, 3 MHz. Scale bar = 50 µm. (i) Pixel distribution of THG signal in the tibia marrow: background (brown rectangle), THG^lo^ (cyan rectangle) and THG^hi^ (yellow rectangle). 2D image of THG signal in cells in the bone marrow, showing THG^lo^ signal in cell membranes (cyan arrowheads) and granular intracellular THG^hi^ signal (rose stars). Scale bar = 30 µm. (ii) Pixel distribution of THG signal in the tibia cortex, with the same color-coding for background, THG^lo^ and THG^hi^ pixels as in the bone marrow. 2D image of THG signal in the bone tissue, showing lacunae and connecting canaliculi. Scale bar = 30 µm. **b**. Cell segmentation distinguishing between THG^lo^ (cyan) and THG^hi^ (yellow) cells (magenta rectangle in **a**). **c**. Merged (left) and single channel (middle and right) 3D image (442×442×102 µm³, 1036×1036×52 voxel) of THG^hi^ (green) and endoplasmic reticulum (ER, magenta) in the bone marrow of an explanted C57/Bl6 mouse tibia (3PM at 1650 nm). Scale bar = 50 µm. **d**. Percentage of ER^+^ cells in the THG^hi^ cell population and of THG^hi^ cells in the ER^+^ cell population (n = 10 mice), showing a strong correlation of THG^hi^ signal and ER staining at the single cell level in the tibia marrow. **e**. Representative close-up images of THG^hi^ER^+^ cells, showing non-identical overlap of THG^hi^ signal and ER staining. **f**. Representative 3D image from a 60 min time-lapse *in vivo* 3PM video at 1650 nm, 3 MHz in the tibia of a CD19:tdRFP mouse (400×400×30 µm³, 518×518×11 voxel), every 30s. Scale bar = 50 µm. **g**. Percentage of THG^hi^ cells in the CD19:tdRFP marrow cell populations with V>500 µm³ (plasma cells) and with V<500 µm³ (B cells), and in the Blimp1:GFP marrow cell population with V>500 µm³ (plasma cells). The percentage of ER^+^THG^hi^ cells in the Blimp1:GFP ER^+^ cell population with V>500 µm³ (plasma cells) is shown in red. **i**. Distributions of mean displacement rates in the THG^hi^ and THG^lo^ plasma cell subset, respectively, showing that THG^hi^ plasma cells are less motile than their THG^lo^ counterparts. Statistical analysis was performed using two-way ANOVA with Bonferroni post-test or two-tail t-test, with p values indicated.

As we assumed highly granular THG^hi^ signal inside cells stems from organelles, rich in lipid bilayers, we investigated whether the endoplasmic reticulum (ER) is abundant in THG^hi^ cells of the bone marrow. Therefore, we labeled the ER in explanted tibia bones of C57/Bl6J mice with ER-Tracker red (BODIPY^TM^ glibenclamide) and performed 3D imaging in the bone marrow at 1650 nm (Fig. 7c). In these 3D images, we segmented THG^hi^ cellular structures with a volume between 65 and 4189 µm³ and counted cells showing abundant ER tracker fluorescence. Additionally, we segmented ER^+^ cells in the same cell volume range and counted those cells displaying THG^hi^ signal. The detection limit of abundant ER tracker fluorescence signal was defined as the highest background count measured within cortical bone areas, in which no ER is expected to be present (Suppl. Fig. 9). We found that 93.1±4.9 % of segmented THG^hi^ cells in the bone marrow display ER tracker fluorescence, and 94.0±3.9 % of the segmented ER^+^ cells display THG^hi^ signal (Fig. 7d). Thus, we concluded that THG^hi^ signal is indicative for abundant ER in bone marrow cells, needed for protein biosynthesis. We expect that THG^lo^ cells also contain ER, however, at a lesser extent than THG^hi^ cells.

Notably, we found that ER tracker and THG^hi^ signal only partially co-localize in the ER^+^ THG^hi^ double positive cells (Fig. 7e). This observation indicates that THG^hi^ signal may originate also from membrane-rich organelles other than endoplasmic reticulum, such as mitochondria. We may not detect the THG^hi^ signal of certain ER structures due to discrepancies between laser polarization and orientation of the lipid bilayers in the ER membrane.

We confirmed the validity of our finding that only ≈1/3 of the plasma cells show THG^hi^ signal in the tibia marrow of CD19:tdRFP mice (Fig. 7f), by analyzing explanted tibia bones of Blimp1:GFP mice (Suppl. Fig. 9). In Blimp1:GFP mice, plasma blasts and plasma cells express GFP, among other cells. The expression level in other cell types is however much lower than in plasma blasts and plasma cells, and their cell volumes are typically lower than those of plasma cells. Hence, we were able to unequivocally identify plasma cells and plasma blasts in the tibia marrow of Blimp1:GFP mice by three-photon 3D imaging at 1330 nm. We used 1330 nm to enable GFP detection upon three-photon excitation. We segmented GFP^+^ cells with a volume between 500 and 4189 µm³, to exclude smaller cells, and analyzed the presence of THG^hi^ signal in these cells. Similar to the data acquired in CD19:tdRFP mice, we found that 36.7±2.7 % of the GFP^+^ plasma cells are THG^hi^.

In some of the explanted tibia bones from the Blimp1:GFP mice we additionally labelled the endoplasmic reticulum using ER tracker and performed 3D three-photon imaging at 1330 nm (Suppl. Fig. 9). Using the previously published SIMI algorithm for spectral unmixing^52,57^ followed by cell segmentation, we found that GFP^+^ THG^hi^ ER^+^ cells represent 34.6±4.5 % of all GFP^+^ cells with a volume between 500 and 4189 µm³ in the tibia marrow. Neither GFP^+^ THG^hi^ cells without ER tracker signal nor GFP^+^ ER^+^ cells without THG^hi^ signal were detected. Thus, THG^hi^ signal is associated with abundant ER also in marrow plasma cells, indicating that THG^hi^ plasma cells have a higher capacity of antibody production, as compared to THG^lo^ plasma cells. Thus, the THG^hi^ signal resolves between two functionally distinct plasma cell subsets in the bone marrow (Fig. 7g).

How the functional state of marrow plasma cells indicated by THG^hi^ signal relates to plasma cell motility can only be addressed by dynamic *in vivo* imaging experiments in the deep marrow of long bones. Our cell tracking analysis of plasma cells in the tibia marrow of CD19:tdRFP mice *in vivo*, at 1650 nm, 3MHz excitation (Fig. 7f, Video 9), showed that plasma cells colocalizing with THG^hi^ signal have a lower mean displacement rate (0.25±0.07 µm/min, n = 31 cells) than THG^lo^ plasma cells (0.44±0.16 µm/min, n = 100 cells), in Fig. 7h (results validated in n = 3 mice). This result indicates that THG^hi^ plasma cells are sessile, whereas THG^lo^ plasma cells are rather migratory.

## Discussion

### Development of dynamic in vivo three-photon imaging in intact tibia

As the birthplace of most hematopoietic cells^9,58^, responsible for immunological memory maintenance^5,6,59^, and being involved in tumor dormancy and metastatic recurrence^14^, the bone marrow microenvironment has been intensively investigated. To give insight into the sequence of events contributing to these long-term processes spatio-temporal studies are of particular importance. Intravital two-photon microscopy (2PM) allowed us to analyze motility patterns of immune cells and their interactions with the environment, in murine marrow of flat bones^21^, of intact long bones of young or irradiated mice^10,11^, after surgical thinning of bone cortex in adult mice^5,15,60^, or by inserting micro-endoscopic lenses into the marrow cavity^24,61^. However, minimally invasive technologies for long bone imaging are still needed.

As there is increasing evidence for functional differences in the immune compartment between the hematopoietic islands in the skull, a flat bone, and the marrow of long bones^17^, dedicated *in vivo* imaging methods are required for the various bone types. The interactions of immune cells with the stromal and vascular compartments have been shown to impact on immune cell functions in the bone marrow^4,8,9^. The need for *in vivo* imaging methods in long bones is supported by the fact that the high mechanical load specific for this bone type, as opposed to flat bones, is expected to have a strong impact on osteo-immune interactions^62^. As in long bones, endosteal areas and deep marrow were found to differ with respect to microenvironmental conditions^4,24,63^, developing technologies which allow dynamic imaging in the deep marrow cavity is key to understand immune cell functions.

Scattering and absorption of near-infrared radiation in two-photon microscopy limit highly resolved, direct optical access to areas located deep within the marrow cavity of intact long bones, through >100 µm thick cortical bone. The straightforward answer to this challenge is the use of long wavelength, infrared radiation at high pulse energy, with narrow fs-pulses, to favor higher order non-linear processes, such as three-photon excitation^31,35^. State-of-the-art optical parametric amplifiers (OPA) have been previously used as excitation sources for three-photon microscopy (3PM), to image various organs and tissues, *in vivo* and *ex vivo* ^29,30,32,35,39,43^, even in freely moving mice^38^. However, *in vivo* 3PM of deep marrow cavity in intact long bones of adult mice, with bone representing the most scattering tissue in living organisms^26^, has not yet been performed, and adequate 3PM setup parameters have not yet been defined for this application. Besides, a main challenge of existing OPA systems for 3PM is imposed by their low repetition rate (0.3 to 2 MHz^29^), leading to slow image acquisition over large fields of view, and, by that, limiting the capacity to investigate dynamic biological processes in the bone marrow, such as immune cell migration. For instance, T cells, the fastest cell population in secondary lymphoid organs, migrate at speeds up to 15 µm/min^64^.

In this work, we demonstrate fast dynamic *in vivo* three-photon imaging of 400×400 µm² tissue areas in the bone marrow of intact mouse tibia, using a novel OPA prototype as 3PM excitation source, which emits high pulse energy (>400 nJ) radiation at 1650 nm and repetition rates in the range 1-4 MHz. The generation of 3 and 4 MHz under the specified conditions is a unique feature of our newly developed OPA prototype. We succeeded to monitor heterogeneous blood flow dynamics and to analyze B lymphocyte motility in the marrow cavity of tibia, through up to 200 µm thick cortical bone, when using 3 and 4 MHz repetition rates, at 20-40 nJ pulse energy at the sample. In benchmarking experiments comparing our 3PM technology to existing 2PM and 3PM methods, we demonstrated that these laser parameters are key to achieve (i) highly resolved imaging as deep as 400 µm in the marrow cavity of intact mouse tibia, (ii) in a time-lapse manner, without any evidence of tissue photodamage neither in bone cortex nor in bone marrow. 400 µm imaging depth in intact tibia corresponds to ≈1.5 fold of the effective attenuation length *l_e_* of radiation in this organ. As imaging depths as high as 10x *l_e_* have been theoretically predicted^65^, we expect that further technological improvements, e.g. adaptive wave front correction, already demonstrated for brain cortex imaging through 100 µm skull bone^36^, will enable imaging throughout the entire volume of long bones in the future. However, the prerequisite for correcting wave front distortions in long bones is that enough laser pulse energy for efficient signal generation arrives at the imaging site deep within tissue, and that emitted fluorescence and higher harmonics signals, typically in the visible spectral range, reach the detector. This is challenging in intact long bones, as both incoming and outgoing radiation is diminished by scattering, especially through thick bone cortex. In the same line, the drawback of 1650 nm radiation able to efficiently excite red fluorescent proteins and dyes, but not broadly used fluorophores, such as GFP, seems to be marginally relevant for intact long bone imaging, as short wavelength fluorescence will hardly succeed to surpass the overlaying bone and marrow tissue layers. A solution to further reduce scattering is the use of even longer excitation wavelengths, and of fluorescent proteins and dyes emitting in the infrared range^66,67^, which call for further laser and microscope optics development^68^, generation of novel infrared fluorescent proteins and of reporter mice based on those.

Reduced scattering at higher excitation wavelengths and the cubic dependence of laser photon flux density for non-resonant three-photon excitation and THG allowed us to achieve highest spatial resolution upon excitation at 1650 nm, in the deep marrow cavity of intact tibias. Considering the expectation of higher diffraction-limited spatial resolution (i.e. at tissue surface) at shorter wavelengths, this appears counterintuitive at first glance^69^. However, it highlights once more the advantage of our OPA prototype to counteract scattering effects in tissue for best imaging performance. The subcellular spatial resolution of 3PM at 1650 nm in deep tissue is underlined by high SNR values. To further reduce noise and, by that, to improve SNR for better cell segmentation and cell tracking in our data, we trained and successfully applied an existing deep-learning algorithm, i.e. Noise2Void^54^, on time lapse 3D fluorescence images of tdRFP^+^ B lineage cells in the tibia marrow. Experimental SNR improvement can be achieved by time-gated detection, as applied in 3PM of the brain cortex^36^. This represents a reliable strategy, as even for higher OPA repetition rates (3 and 4 MHz), noise detection temporally surpasses the detection of fluorescence and harmonics generation signals by several orders of magnitudes. Still, time gated detection requires customized modifications of the microscope, which are not readily available for a broad application, in contrast to the here proposed de-noising pipeline. Besides, for the detection of phosphorescence, e.g. when using oxygen-sensitive probes^70^ to monitor tissue oxygenation *in vivo*^71^, time-gating is no longer favorable, as phosphorescence lifetimes are in the same range as the time window between two consecutive laser pulses of OPA systems.

The downside of using high pulse-energy long-wavelength radiation for *in vivo* imaging is the inherent danger to damage the tissue. By performing immunofluorescence analysis, we have shown that non-resonant three-photon excitation at 1650 nm, while potentially expected to overheat biological samples, is not associated with tissue thermal or photodamage. We verified this finding by monitoring intact blood flow *in vivo*, in the tibia marrow at 1650 nm, 3 and 4 MHz. Along the same line, we found that marrow B lymphocyte mean velocity and displacement rate values measured by 3PM at 1650 nm were similar to those measured by 2PM at 1100 nm, in the same tissue area.

No signs of damage were detected at the bone surface above the imaged marrow sites. Finally, photobleaching of tdRFP, the red fluorescent protein used for time-lapse *in vivo* imaging, was negligible confirming the reliability of the here described 3PM method.

### 3PM and label-free THG in vivo imaging provides new insights into marrow biology and links heterogeneous functional capacity to migratory behavior of plasma cells

*In vivo* label-free THG imaging using our 3PM method highlights the 3D tissue architecture of both bone and marrow compartments in the intact tibia, with high fidelity, at subcellular resolution. Performed in a time-lapse manner, we could show that THG imaging enables monitoring whole-tissue dynamics, and may represent a powerful tool to monitor mechanical cues, such as shear stress in vessels and tissue strain, both in stiff and soft tissues. Prospectively, the THG signal will be able to inform about the 3D force field acting on single immune cells *in vivo*, with impact on their functions^62^.

Moreover, the in-depth analysis of THG signal in the tibia marrow allowed us to reliably associate THG^hi^ signal in cells with a high organelle content distinguishing these cells from THG^lo^ cells, in which the THG signal mainly originates from the cell membrane, in line with previous reports on isolated leukocytes^44^. As we found THG^hi^ signal to be unequivocally associated with abundant endoplasmic reticulum (ER) in all marrow cells, this label-free, ubiquitous signal has the potential to indicate cellular functional states associated with protein biosynthesis in any cell type, if co-registered with cell type specific labelling, e.g. in fluorescent reporter mice.

Analysis of *in vivo* 3PM data acquired in the unperturbed tibia marrow of B lineage reporter mice allowed us to show that THG^hi^ signal is enriched in plasma cells as compared to B cells. However, only ≈1/3 of marrow plasma cells were THG^hi^, shown also to have abundant ER. As ER is required to produce large amounts of antibodies, we conclude that THG^hi^ signal defines heterogeneity of functional capacity among marrow plasma cells *in vivo*. In order to account for their antibody producing function, B cells massively expand their ER when differentiating into plasma cells^72^.

Our 3PM method enables intravital time-lapse imaging of currently inaccessible marrow regions in intact tibia of B lineage reporter mice, over large fields-of-view. Hence, we could analyze the migration behavior of marrow plasma cells over up to three hours, in a statistically reliable manner, as we could monitor >100 plasma cells per animal. In line with previous reports^5^, we found two plasma cell subsets with distinct motility patterns, i.e. a non-migratory and a highly motile subset, characterized by mean displacement rates of ≈0.2 µm/min and ≈1 µm/min, respectively.

By co-registering THG signal with plasma cell labeling in the time-lapse 3PM data, we found that THG^hi^ plasma cells are non-migratory, with a mean displacement rate of ≈0.25 µm/min, whereas highly motile cells are THG^lo^ plasma cells, with a mean displacement rate of ≈0.45 µm/min. These data suggest that the capacity of plasma cells to produce large amounts of antibodies is inversely linked to their migratory behavior. Plasma cell residing in their survival niches are non-migratory^5,8^, and are able to produce large amounts of antibodies, as in these niches they find suitable microenvironmental conditions supporting their metabolic demands^71^. In contrast, migratory plasma cells, including plasma blasts as precursors of sessile plasma cells^73^, presumably searching for an appropriate microenvironment, may have a limited capacity for protein biosynthesis, as previous reports showed varying antibody secretion by plasma cells dependent on extrinsic factors in tissue^74^.

Taken together, in this study, we developed a laser with unique 3 and 4 MHz repetition rates of high-pulse-energy 1650 nm radiation and used it as excitation source for dynamic *in vivo* three-photon imaging of intact mouse tibia. In this way, we were able to access the deep marrow cavity in a minimally invasive manner and, together with in-depth analysis of ubiquitous label-free THG signal and cell type specific fluorescence, to investigate links between cellular motility patterns and functional capacity related to protein synthesis, opening unprecedented opportunities to understand bone biology *in vivo*.

## Material and methods

### Experimental setup for multi-photon tibia imaging

#### Excitation sources

The excitation sources for combined two- and three-photon microscope are two optical parametric amplifiers (OPA), typically using chirped laser pulses to support high pulse energies: (1) novel OPA design (Ytterbia OPA), Thorlabs Inc. Lafayette CO, USA, with a Ytterbium-fiber based pump laser, 1030 nm, 20 W and (2) tunable AVUS OPA (APE GmbH, Berlin, Germany) pumped by Aeropulse FS20 (NKT Photonics, Birkerod, Denmark). Pulse compression was achieved using pre-chirp ZnSe plates (for Ytterbia OPA) and a fixed design of a two-prism pulse compressor (for AVUS OPA). The emission wavelength of the Ytterbia OPA is fixed at 1650 nm, bandwidth 60 nm, with tunable repetition rate between 1.01 MHz and 3.98 MHz. The AVUS OPA systems operates in the wavelength range from 1200 nm to 2500 nm (idler), at fixed repetition rate of 2 MHz. The Ytterbia OPA at 1650 nm provides an average power of 1.8 W (452 nJ) at 3.98 MHz, 1.49 W (481 nJ) 3.09 MHz, 1 W (486 nJ) at 2.06 MHz and 0.5 W (495 nJ) at 1.01 MHz. The AVUS provides an output average power at 1330 nm of 700 mW (350 nJ per pulse at 2 MHz repetition rate) and at 1630 nm of 500 mW (250 nJ per pulse at 2 MHz repetition rate). We measured the pulse duration both directly at the laser output and under the objective, by second-order interferometric autocorrelation using CARPE (coating for spectral range 1200 nm - 1700 nm, APE GmbH, Berlin, Germany). The combination of λ/2 waveplate and beam-splitter cube polarizer is used to control laser power. The excitation sources for state-of-the-art two-photon microscopy are a mode-locked Titanium-Sapphire laser (Ti:Sa, 690-1080nm, 80 MHz Chameleon Ultra II, Coherent, Glasgow, UK) tuned at 930 nm and an optical parametric oscillator (OPO, 1050-1350 nm, 80 MHz, APE GmbH, Berlin, Germany, pumped by the Ti:Sa) tuned either at 1100 nm or at 1330 nm. A high-accuracy positioning mirror is used to switch between the OPA and OPO laser beams. The Ti:Sa and optical parametric beams are combined by a dichroic mirror and overlapped in the microscope scan head.

#### Microscope setup

The laser-scanning microscope used in this study is based on a commercial two-photon microscopy system (TriMScope II, LaVision BioTec, Bielefeld, Germany)^52^, in which we replaced the optics for broad spectral transmission up to 1700 nm, matching the polarization of all excitation sources. Water-immersion objective lenses (Olympus, XLPLN25XWMP2, 25x, NA 1.05; Nikon, CFI75 Apo 25XC W 1300, 25x, NA 1.1, Tokyo, Japan) are used to focus the excitation laser beams into the sample. Fluorescence, SHG and THG signals are collected through the objective lenses in the epi-direction using a dichroic mirror (900, Chroma, US) and directed towards four photomultiplier tubes (PMTs, H7422-40/-50, Hamamatsu, Japan). All PMTs are assembled in a detection system with four optical channels, where each channel is defined by individual interference filters and a set of dichroic mirrors. The assignment of optical channels depends on the excitation wavelength. The optical filters for excitation at 930 nm (Ti:Sa) are 447±20 nm for SHG, 525±25 nm for the GFP signal, 595±25 nm for the tdRFP and tdTomato signals, at 1100 nm (OPO), 562±20 nm for the SHG signal and 595±25 nm for the tdRFP and tdTomato signals. At 1330 nm (OPA or OPO), following filters were used: 447±20 nm for THG, 525±25 nm for the GFP fluorescence, 595±20 nm for the tdTomato or ER tracker fluorescence and 655±20 nm for the SHG signal. At 1630 nm/ 1650 nm (of both OPA systems), we used the following filters: 562±20 nm for THG signal, 595±20 nm for the tdTomato, tdRFP or ER tracker fluorescence and 810±45nm for the SHG signal. Acquisition time, pixel dwell time and z-step for the 3D z-stacks are stated in the *Results* and *Figure legends*. Time-lapse 2D image acquisition was performed at 1 Hz (every 1 s) and 2 Hz (every 0.5 s). Time-lapse 3D image acquisition was performed every 30 s over up to 1 hour or every 120 s over 2 hours.

### *Ex vivo* NanoCT imaging

The explanted tibiae were placed in a hollow plastic cylinder filled with Styrofoam to prevent movement. The cylinder was glued to a glass rod and fixed in the sample holder of the nanofocus-CT system (Phoenix Nanotom M, GE Sensing & Inspection Technologies GmbH, Wunstorf, Germany). CT scans were acquired at a voltage of 100 kV, a current of 120 µA, and a 0.1 mm copper prefilter. One measurement included 1500 radiograms (3 scans per angular step for noise reduction, with 500 ms acquisition time per single scan), the total acquisition time being 54 minutes. The magnification was 50x, with a voxel edge length of 2.00 µm. The scans were reconstructed as 3D volumes using the phoenix datos|x-ray2 reconstruction 2.4.0-RTM software (GE Sensing & Inspection Technologies GmbH, Wunstorf, Germany).

### Scanning electron microscopy of explanted tibia

The explanted tibiae were fixed in 1% paraformaldehyde (PFA) and dehydrated in a series of exchanges with increasing concentrations of ethanol (99.8%) diluted in distilled water (50%, 70%, 80%, 90%, 95%) for 4 minutes each. Finally, they were kept in 100% concentration ethanol for 20 minutes, followed by 5 rinses in liquid carbon dioxide, at the critical draying point of CO_2_ (31.1° C at 73 atm). The dehydrated specimens were stored in phosphate buffer solution (PBS). The samples were fixed with conductive adhesive on sample holders and sputtered with 25 nm platinum. Imaging was performed with a Zeiss ULTRA Plus scanning electron microscope (Carl Zeiss NTS GmbH, Jena, Germany) at an electron high tension of 5.00 kV, with an aperture size of 30.00 µm and a working distance of 11.2 mm to 13.2 mm. Magnification in the shown imaged was in a range of 24x to 26x.

### Image processing and analysis

#### Spectral unmixing of multi-photon images using similarity unmixing (SIMI)

We previously developed similarity unmixing algorithm (SIMI), a numerical pixel-based algorithm, to resolve mixtures of cellular and tissue compartments labeled by different fluorophores or non-linear harmonic generation signals, such as SHG and THG, using a low number of detection channels^52,57^. In the conventional SIMI pipeline, fingerprints of individual cellular and tissue compartments are determined from separate single-color measurements. In the multicolor 3D images acquired in the tibia of Blimp1:GFP mice after ER staining at 1330 nm, we expected overlapping signals in single elementary voxels, i.e. THG signal, GFP fluorescence and fluorescence of endoplasmic reticulum (ER) staining may appear together within one voxel. Thus, we needed to extend the initial SIMI algorithm and to identify *in situ* fingerprints in the analyzed multiplexed images, generated using four detection channels (447±20 nm, 525±25 nm, 595±20 nm and 655±20 nm). We found and assigned fingerprints to three mixed components, THG+ER (bone marrow), THG+GFP+ER (bone marrow) and THG+SHG (bone cortex), and to two individual signals, GFP and THG (Suppl. Fig. 9). SIMI unmixing was performed based on similarity analysis, i.e. searching for the closest match between the pixel intensities of the raw image and the signal intensity of the determined fingerprint for each detection channel.

### Image denoising using the Noise2Void deep-learning algorithm

To improve SNR by reducing the noise level in the time-lapse 3D images of tdRFP^+^ B lineage cells in the tibial bone marrow, we used a machine-learning algorithm, Noise2Void (N2V)^54^ plug-in as a part of the open-source neural network algorithms CSBDeep^75^ package in FIJI. The trainable deep-learning scheme of N2V plug-in allows to predict the original shape of the cells without clean targets. This was used to improve cell segmentation and tracking. The fluorescence signal in each pixel is restored by reducing the estimated noise, which distribution differs in a certain neighborhood, known as the receptive field. The training of the N2V-models was performed on individual 2D images from different time points and tissue depths, in different experiments, to enhance model generalization. To account for PMT properties, we performed training on each detection channel individually. The model setting was 20 images for each run, 200 epochs, 300 steps per epoch, 128/256 batch size per step, and neighborhood radius of 5.

### Cell and tissue segmentation, cell tracking

#### Cell segmentation and classification

3D reconstruction of z-stacks and NanoCT images was performed in Imaris (software version 9.7.2, Bitplane, an Oxford Instruments company, Belfast, UK). Cell segmentation in multi-photon 3D stacks and time-lapse 3D images was automatically performed using Imaris, after machine-learning-based noise reduction using the trained N2V algorithm in FIJI, version 1.53t as previously described. Motion correction of the time-lapse 3D images in Video 7 was performed using the Fast4Dreg plugin^76^ in FIJI, before further processing the data. For object surface detection in Imaris, we used background subtraction relying on local contrast and separated touching objects, relying on seed point diameter of 7.5 µm. Co-localization with co-registered additional signals in both 3D stacks and time-lapse 3D images was performed using the filtering function in Imaris, after cells were segmented. The same function was used to classify segmented cells by volume, e.g. to distinguish between B and plasma cells in CD19:tdRFP mice.

#### Cell tracking

Automatized cell tracking in time-lapse 3D images was performed using Imaris, after cell segmentation. Therefore, we used the autoregressive motion algorithm in Imaris, with a maximum frame-to-frame distance of 8 µm and maximum number of gaps = 3. We verified all tracking results and, in some cases, disconnected tracks of one and the same cell were manually connected.

#### Cortical thickness

The cortical thickness of the tibia was determined by applying the FIJI function *Local Thickness* on the xz- or yz-resliced 3D images of SHG and THG signals in the tibia.

### Animal procedures

The CD19:tdRFP B lymphocyte fate mapping mice^11^, Prx1:tdRFP mesenchymal stromal cells fate mapping mice^77^ and Cdh5:tdTomato x Histone:GFP (Cdh5:tdTom) fate mapping mice^61,78^ (kind gift of Prof. R. Adams, Dr. M.G. Bixel) used for *in vivo* imaging were typically 12-24 weeks old females and males. Additionally, explanted tibiae from old Cdh5:tdTom mice (106 weeks and two 127 weeks old female mice), from Blimp1:GFP mice and from C57/Bl6J mice were used in this study. All animal experiments were conducted in accordance with the ARRIVE guidelines and approved by Landesamt für Gesundheit und Soziales, Berlin, Germany in accordance with institutional, state and federal guidelines (G0048/21 and G0158/16). In intravital experiments, the right tibia of n = 12 CD19:tdRFP mice, n = 16 Cdh5:tdTom mice and n = 2 Prx1:tdRFP mice, and the spleen of n = 3 CD19:tdRFP mice were imaged. Explanted tibias from n = 3 Cdh5:tdTom mice, from n = 6 Blimp1:GFP mice and n = 10 C57/Bl6J mice were imaged.

### Intravital surgery for imaging

All surgery preparations for intravital imaging were initialized with the following steps. The mice were placed on a heating plate at 37°C and anesthetized by inhalation narcosis with isoflurane, corresponding to their weight.

#### Tibia preparation

The right hind paw was fixed, by stretching the leg to the side, and prepared for further surgery. To prevent contamination, the fur was shaved clean on the upper side of the fixed paw. In the diaphysis area of the tibia, the skin layer was dissected, and the muscle fibers were pushed aside to get access to the flat, medial surface of the tibia shaft. The tibia was kept in position by fixing the crest with a surgery clamp. A wall of 1 % agarose gel was built around the opened area to form a cylindrical bath filled with isotonic NaCl solution for imaging. The medial part of the tibia was imaged, as indicated in Fig. 1(iv). Alternatively, tibia cortex was additionally thinned with a diamond drill, for comparative imaging at low vs. high-pulse energy (2PM vs. 3PM).

#### Spleen preparation

The mice were positioned on the right side to get access to the spleen from the top. To keep the dissection area clean, the hair was removed in the location above the spleen. Through a small excision in the skin a part of the spleen was pulled out and placed on a heating plate, which was kept at 37°C. The opened space between the part of the spleen and the dissection area was sealed with glue. The agarose bath filled with isotonic NaCl solution was constructed around the spleen part exposed for intravital imaging.

### Explanted tibia preparation for imaging

C57/Bl6J or *Blimp1:GFP* mice were sacrificed by cervical dislocation. Tibia bones were explanted and muscle and connective tissues were removed, tibia was glued on a Petri dish, which was then filled with PBS and immediately imaged.

### Endoplasmic reticulum staining in explanted tibia

The freshly explanted tibiae were placed in a Petri dish containing ER Tracker^TM^ Red (BODYPI^TM^ TR Glibenclamide) for cell imaging (Invitrogen, Eugene, Oregon, USA) in PBS medium), at 37°C for 30 minutes. For that, 5 µl stock solution of ER Tracker^TM^ Red (MW 915.23 g/mol) in DMSO (concentration 1 mmol/l) were diluted in 5 ml PBS. The epiphyses of the bones were removed for a better dye penetration. The tibiae were washed with PBS to remove exceeding dye and glued on a Petri dish, containing PBS, for imaging purposes.

### Tissue preparation for histology

Freshly explanted tibiae and spleens were fixed in 4% PFA for 4 hours, cryoprotected in 10-30% sucrose/PBS, embedded in O.C.T. and stored at −80°C. 7-micrometer-thick serial spleen sections were cut using a cryostat and collected on positively charged slides. Fixed tibiae were covered with Kawamoto’s medium (SCEM, Section-Lab Co. Ltd., Hiroshima, Japan), frozen and cryo-sectioned using the Kawamoto’s film method^79^. Sections were kept at −80°C until further use.

### Immunofluorescence and Movat’s pentachrome histological analysis

For immunofluorescence, slides were dried, washed in PBS, blocked with 10% serum, and stained with antibodies in PBS, containing 1% serum and DAPI, for 1–2 h. The following antibodies were used: Ly6G (PE-conjugated), CD68 (AF647-conjugated, FA-11, Biolegend), CD45R (B220) antibody (FITC-conjugated, anti-mouse REAfinity, Miltenyi GmbH, Bergisch Gladbach, Germany), CD3 antibody (FITC-conjugated, Biolegend), HSP70 (unconjugated, 4872, Cell signaling), anti-rabbit (AF647-conjugated). Stained slides were mounted with Fluoromount mounting medium (Thermo Fisher, MA, US). TUNEL staining was performed using an ApopTag Fluorescein *in situ* apoptosis detection kit (S7110, EMD Millipore, Germany) according to the manufacturer’s protocol. Briefly, the O.C.T.-embedded sections were thawed and rehydrated, and then permeabilized in cooled acetic acid and in ethanol for 3 min at −20°C. Sections were incubated with the reaction buffer containing TdT enzyme at 37°C for 1 h. After washing with stop/wash buffer, sections were treated with anti-digoxigenin-FITC for 30 min at room temperature. The sections were counterstained with DAPI before being mounted with Fluoromount mounting medium. Alternatively, 7 µm tibia sections were stained using the Movat’s pentachrome staining^80^. Both immunofluorescence and Movat’s pentachrome staining histological analysis was performed on a fluorescence wide-field microscope (Keyence BZ800, Keyence Deutschland GmbH, Neu Isenburg, Germany), using magnifications of 4x, 10x, and 20x.

### Point-spread function measurement

The spatial resolution of our imaging setup was determined as the lateral and axial dimensions of the effective point spread function (ePSF), measured relying on the highly-resolved 3D fluorescence signal of 200 nm size nanospheres (605 nm emission wavelength, F8810 FluoSpheres, Invitrogen, Eugene, Oregon, USA), embedded in agarose gel (1%). The nanosphere are used as a standard for PSF measurements, as their size is below the resolution limit of the imaging setup. Following the Nyquist criterion, the voxel size for the PSF measurements was 0.1 µm laterally (xy) and 0.5 µm axially (z). x/y and z fluorescence intensity profiles of the nanospheres were plotted and approximated by Gaussian (in FIJI, confirmed by fitting in Origin 10.0) to determine the dimension of the ePSF in the respective direction.

### Statistical analysis

Statistical analysis was performed using Origin 2023 (OriginLab, CA, US) or GraphPad Prism 7.0c (GraphPad Software Inc., San Diego, CA, US).

## Supporting information

Complete supplemental information, including supplemental figures.

## Authors contributions

A.R. built, optimized and characterized the 2PM/3PM imaging system. A.R., A.F.F, R.G., R.A.N., R.L., Y.C performed the intravital experiments and imaging experiments on explanted tissues. J.H. and V.A. helped to optimize the imaging system and S.D. and L.W. developed the Ytterbium-based OPA laser together with A.R and R.A.N. A.E.H. and C.U. provided expertise in immune dynamics, plasma cell biology and regarding all animal experiments. Y.C. provided expertise regarding ER staining in long bones. J.R. and A.E.H. designed and performed the immunofluorescence experiments concerning the possible tissue photodamage by 3PM. R.K. trained the N2V deep-learning algorithm. R.K. and A.F.F. performed image denoising. G.D. provided key expertise concerning quantification of bone surface damage upon 3PM. A.B. and A.H. performed the Nanofocus-CT experiments. A.B. and I.B. performed the scanning electron microscopy experiments. A.R., A.F.F., A.E.H. and R.A.N. analyzed data, processed the results, and prepared the figures. A.R., A.F.F., A.E.H. and R.A.N. wrote the manuscript. A.E.H. and R.A.N. supervised the study. All authors edited the manuscript.

## Acknowledgments

The research leading to these results has received funding from the Deutsche Forschungsgemeinschaft (DFG), Germany under grant CRC 1444, P14 to R.A.N. and A.E.H. and P09, P13 to G.D., under grant FOR 5560, P02 to R.A.N (NI1167/9-1) and A.E.H (HA5354/13-1), under grant HA5354/12-1 to A.E.H. and under grant 317850156 to A.H. Additionally, funding from the Einstein Research Foundation, Berlin under grant ESB-A-2019-559 to A.E.H. and R.A.N. The authors thank R. Uecker, P. Mex and G. Korus for excellent technical assistance. The authors thank D. Walgurski (Helmholtz-Zentrum Berlin) for support in performing electron microscopy studies.

## Data availability statement

The data that support the findings of this study are available on the data repository Zenodo, under the links: https://doi.org/10.5281/zenodo.8383833 and https://doi.org/10.5281/zenodo.7464124.

## Code availability statement

Customized code for SIMI analysis, rose plot generation and the trained N2V algorithm are provided by the authors.

