## Supplementary material for "*In vivo* optimized three-photon imaging of intact mouse tibia links plasma cell motility to functional states in the bone marrow": Complete supplemental information, including supplemental figures.

### 1. Theoretical background of multi-photon excitation processes and signal analysis in multi-photon microscopy

#### 1.1. Fluorescence signal upon non-resonant or resonance-enhanced two- and three-photon excitation

We consider that a fluorophore molecule transits only virtual energetic states during a non-resonant excitation, either two- or three-photon excitation. Only the initial and final energetic states of the fluorophore molecule are real energetic states. During a resonance-enhanced three-photon excitation, the molecule also transits real energetic states, not only virtual states, which increases the excitation probability and, by that, leads to an increased excitation rate as compared to a non-resonant three-photon excitation process.

The experimentally measured fluorescence signal is given by the fluorescence signal per laser pulse *F*, integrated over the pixel dwell time, i.e. the number of excitation pulses within the pixel dwell time.

The fluorescence photon flux density per laser pulse *F*, originating from a sample containing molecules of a single fluorophore type, is given by[1-4]:

$$F_{2P} = \frac{1}{2} {N_{abs}}^{2P} N_{abs}^{2P} \int_0^{\infty} \varphi(\lambda_{em}) \cdot \zeta(\lambda_{em}) d\lambda_{em} \tag{1}$$

for a non-resonant two-photon excitation process and

$$F_{3P} = \frac{1}{3} {N_{abs}}^{3P} \int_0^{\infty} \varphi(\lambda_{em}) \cdot \zeta(\lambda_{em}) d\lambda_{em} \tag{2}$$

for a non-resonant three-photon excitation process. In Eq. (1) and (2) $N_{abs}$ represents the number of absorbed photons, $\varphi$ the fluorescence quantum yield and $\zeta$ the detector quantum efficiency of the experimental setup.

As fluorescence quantum yield – the spontaneous emission of photons by an excited molecule – is considered to be independent of the excitation process and as fluorescence detection in all our experiments does not change, we simplify the integrals in Eq. (1) and Eq. (2) by $\bar{\varphi}$.

$$\bar{\varphi} = \int_0^{\infty} \varphi(\lambda_{em}) \cdot \zeta(\lambda_{em}) d\lambda_{em} \tag{3}$$

The number of absorbed photons per molecule $N_{abs}$ depends on the excitation rate, i.e. either non-resonant two- or three-photon excitation. The excitation rate depends on the capacity of the molecule to simultaneously be excited by two and three photons, respectively. The measure of this capacity is given by the active two-photon absorption cross-section $\sigma_{2P} \cdot \eta_{2P}$ for a non-resonant two-photon excitation process and the active three-photon absorption cross-section $\sigma_{2P} \cdot \eta_{3P}$ for a non-resonant three-photon excitation process. Further, the excitation rate depends on the excitation photon flux density of the laser under the objective lens, i.e. time-dependent intensity of the excitation source. $N_{abs}$ is also dependent on the number of molecules within the excitation volume, given by the concentration *C*. Thus, according to[1,3], the number of absorbed photons is for a two-photon excitation:

$${N_{abs}}^{2P} = C \cdot \sigma_{2P} \cdot \eta_{2P} \cdot \int_t \int_{V \to \infty} S^2(\boldsymbol{r}) \cdot {I_0}^2(t) dt dV \tag{4}$$

and for a three-photon excitation:

$${N_{abs}}^{3P} = C \cdot \sigma_{3P} \cdot \eta_{3P} \cdot \int_t \int_{V \to \infty} S^3(\boldsymbol{r}) \cdot {I_0}^3(t) dt dV \tag{5}$$

with $I_0(t)$ the time dependent laser photon flux and $S(\boldsymbol{r})$ its unitless three-dimensional spatial distribution function.

As temporal and spatial distributions of the excitation photon flux can be considered to be independent from each other, Eq. (4) turns into:

$$N_{abs}{}^{2P} = C \cdot \sigma_{2P} \cdot \eta_{2P} \cdot \int_t I_0{}^2(t)dt \int_{V\to\infty} S^2(r) \cdot dV \qquad (6).$$

If $g^{(2)}$ is the degree of second-order coherence of the excitation source being defined as $\frac{\langle I_0{}^2(t)\rangle}{\langle I_0(t)\rangle^2}$ [5], Eq. (6) becomes:

$$N_{abs}{}^{2P} = C \cdot \sigma_{2P} \cdot \eta_{2P} \cdot g^{(2)}\langle I_0\rangle^2 \int_{V\to\infty} S^2(r) \cdot dV \qquad (7),$$

with ⟨ ⟩ indicating the average over time.

Similarly, Eq. (5) turns into:

$$N_{abs}{}^{3P} = C \cdot \sigma_{3P} \cdot \eta_{3P} \cdot \int_t I_0{}^3(t)dt \int_{V\to\infty} S^3(r) \cdot dV \qquad (8).$$

If $g^{(3)}$ is now the degree of third-order coherence of the excitation source being defined as $\frac{\langle I_0{}^3(t)\rangle}{\langle I_0(t)\rangle^3}$, Eq. (8) becomes:

$$N_{abs}{}^{3P} = C \cdot \sigma_{3P} \cdot \eta_{3P} \cdot g^{(3)}\langle I_0\rangle^3 \int_{V\to\infty} S^3(r) \cdot dV \qquad (9).$$

For solving the temporal integrals $\int_t I_0{}^2(t)dt$ and $\int_t I_0{}^3(t)dt$ , we assume, based on our experimental data, Gaussian laser pulses for both optical parametric oscillator as a two-photon excitation source (OPO) and both optical parametric amplifiers as two and three-photon excitation sources (OPA). According to[2], we define the origin of the time axis at the maximum of the laser pulse, with the pulse width $\tau_p{}^{2P}$ and $\tau_p{}^{3P}$ as well as the repetition rates $RR_{2P}$ and $RR_{3P}$ for the OPO and OPA lasers, respectively. Hence, the temporal integral in Eq(6) for a two-photon excitation becomes:

$$\int_{-1/(2\cdot RR_{2P})}^{1/(2\cdot RR_{2P})} I_0{}^2(t)dt = \frac{g_p{}^{2P}}{\tau_p{}^{2P}} \cdot \left[\int_{-1/(2\cdot RR_{2P})}^{1/(2\cdot RR_{2P})} I_0(t)dt\right]^2 \qquad (10)$$

with $g_P{}^{2P}$=0.664 for a Gaussian beam profile of a single-mode laser pulse ($TEM_{00}$). Thus, the degree of second-order coherence of the excitation source is $g^{(2)} = \frac{g_P{}^{2P}}{RR_{2P}\cdot\tau_p{}^{2P}}$ .

Similarly, the temporal integral in Eq. (8) for a three-photon excitation becomes:

$$\int_{-1/(2\cdot RR_{3P})}^{1/(2\cdot RR_{3P})} I_0{}^3(t)dt = \frac{g_p{}^{3P}}{(\tau_p{}^{3P})^2} \cdot \left[\int_{-1/(2\cdot RR_{3P})}^{1/(2\cdot RR_{3P})} I_0(t)dt\right]^3 \qquad (11)$$

with $g_P{}^{3P}$=0.509 for a Gaussian laser pulse[3]. Thus, the degree of third-order coherence of the excitation source is $g^{(3)} = \frac{g_P{}^{3P}}{RR_{3P}{}^2 \cdot (\tau_p{}^{3P})^2}$ .

Assuming the laser beam is diffraction limited, with *NA* the numerical aperture, i.e. $NA = n \cdot \sin(\alpha)$ with *n* being the refractive index of the medium and 2α the maximal angle of the light cone of the objective lens), the relation between the time-dependent average intensity $\langle I_0 \rangle$ at the focal plane and average laser power $\langle P \rangle$ under the objective lens is defined as[2]:

$$\langle I_0 \rangle = \frac{\pi \cdot NA^2}{\lambda_{exc}{}^2} \cdot \langle P \rangle \qquad (12),$$

with $\lambda_{exc}$ the excitation wavelength.

From Eq. (7), (10) and (12) for a two-photon excitation process, it follows:

$$N_{abs}{}^{2P} = C \cdot \sigma_{2P} \cdot \eta_{2P} \cdot \frac{g_P{}^{2P}}{RR_{2P} \cdot \tau_p{}^{2P}} \frac{\pi^2 \cdot NA^4}{\lambda_{exc}{}^4} \cdot \langle P \rangle^2 \int_{V\to\infty} S^2(r) \cdot dV \qquad (13).$$

Similarly, from Eq. (9), (11) and (12) for a three-photon excitation process, it follows:

$$N_{abs}{}^{3P} = C \cdot \sigma_{3P} \cdot \eta_{3P} \cdot \frac{g_P{}^{3P}}{RR_{3P}{}^2 \cdot (\tau_p{}^{3P})^2} \frac{\pi^3 \cdot NA^6}{\lambda_{exc}{}^6} \cdot \langle P \rangle^3 \int_{V\to\infty} S^3(r) \cdot dV \qquad (14).$$

In the following we use the previously defined[6] unitless coordinate $\nu$ in planes perpendicular to the optical axis, defined as $\nu = \frac{2\pi \cdot NA \cdot \rho}{\lambda_{exc}}$, and the unitless coordinate *u* along the optical axis of the microscope, defined as $u = \frac{2\pi \cdot NA^2 \cdot z}{n \cdot \lambda_{exc}}$, with ρ and z the coordinates in the metric unit system. The spatial distribution of the excitation photon flux under the objective lens is defined as:

$$S(\boldsymbol{r}) = h^2(u, \nu) \qquad (15)$$

with *h(u,ν)* the point-spread function (PSF), which we consider to be Gaussian for both u and ν coordinates based on our experimental observations. Consequently, the spatial integrals $\int_{V\to\infty} S^2(r) \cdot dV$ in Eq. (13) and $\int_{V\to\infty} S^3(r) \cdot dV$ in Eq(14) are estimated to be:

$$\int_{V\to\infty} S^2(r) \cdot dV = \int\int h^4(u, \nu) \cdot du d\nu \approx \frac{n \cdot \lambda_{exc}{}^3}{\pi \cdot NA^4} \qquad (16)$$

for two-photon excitation, in accordance with[2], and

$$\int_{V\to\infty} S^3(r) \cdot dV = \int\int h^6(u, \nu) \cdot du d\nu \approx \frac{2\sqrt{2}}{3\sqrt{3}} \cdot \frac{n \cdot \lambda_{exc}{}^3}{\pi \cdot NA^4} = 0.544 \cdot \frac{n \cdot \lambda_{exc}{}^3}{\pi \cdot NA^4} \qquad (17)$$

for three-photon excitation.

Replacing the integral in Eq. (13) with the result of Eq. (16) and that in Eq. (14) with the result from Eq. (17), we obtain the number of absorbed photons per laser pulse for a two- and three-photon excitation process, respectively:

$$N_{abs}{}^{2P} = C \cdot \sigma_{2P} \cdot \eta_{2P} \cdot \frac{g_P{}^{2P}}{RR_{2P} \cdot \tau_p{}^{2P}} \frac{\pi \cdot n}{\lambda_{exc}} \cdot \langle P \rangle^2 \qquad (18)$$

$$N_{abs}{}^{3P} = C \cdot \sigma_{3P} \cdot \eta_{3P} \cdot \frac{g_P{}^{3P}}{RR_{3P}{}^2 \cdot \left(\tau_p{}^{3P}\right)^2} \cdot 0.544 \cdot \frac{\pi^2 \cdot n \cdot NA^2}{\lambda_{exc}{}^3} \cdot \langle P \rangle^3 \qquad (19)$$

Hence, the fluorescence signal per laser pulse for the two excitation schemes is given by:

$$F_{2P}(t) = \frac{1}{2} \bar{\varphi} \cdot \sigma_{2P} \cdot \eta_{2P} \cdot C \cdot \frac{g_P{}^{2P}}{RR_{2P} \cdot \tau_p{}^{2P}} \frac{\pi \cdot n}{\lambda_{exc}} \cdot \langle P \rangle^2 \qquad (20)$$

and

$$F_{3P}(t) = \frac{1}{3} \bar{\varphi} \cdot \sigma_{3P} \cdot \eta_{3P} \cdot C \cdot \frac{g_P{}^{3P}}{RR_{3P}{}^2 \cdot \left(\tau_p{}^{3P}\right)^2} \cdot 0.544 \cdot \frac{\pi^2 \cdot n \cdot NA^2}{\lambda_{exc}{}^3} \cdot \langle P \rangle^3 \qquad (21).$$

In order to estimate the active three-photon excitation cross-section from the known value of the active two-photon excitation cross-section for the same fluorophore, at the same site of the sample, we use the ratio between the fluorescence signal per laser pulse upon two- and three-photon excitation:

$$\frac{F_{2P}}{F_{3P}} = \frac{\frac{1}{2} \cdot \sigma_{2P} \cdot \eta_{2P} \cdot \frac{0.664}{RR_{2P} \cdot \tau_p{}^{2P}} \frac{1}{\lambda_{exc}{}^{2P}} \cdot \langle P^{2P} \rangle^2}{\frac{1}{3} \cdot \sigma_{3P} \cdot \eta_{3P} \cdot \frac{0.509}{RR_{3P}{}^2 \cdot \left(\tau_p{}^{3P}\right)^2} \cdot 0.544 \cdot \frac{\pi \cdot NA^2}{\left(\lambda_{exc}{}^{3P}\right)^3} \cdot \langle P^{3P} \rangle^3} \qquad (22).$$

By substitution, the active three-photon excitation cross-section $\eta_{3P}\sigma_{3P}$ with the unit $cm^6 \cdot s^2$ is given by:

$$\sigma_{3P} \cdot \eta_{3P} = \frac{3}{2} \cdot \frac{F_{3P}}{F_{2P}} \cdot \sigma_{2P} \cdot \eta_{2P} \cdot \frac{\left(RR_{3P} \cdot \tau_p{}^{3P}\right)^2}{RR_{2P} \cdot \tau_p{}^{2P}} \cdot \frac{\left(\lambda_{exc}{}^{3P}\right)^3}{\pi \cdot NA^2 \cdot \lambda_{exc}{}^{2P}} \cdot 2.398 \cdot \frac{\langle P^{2P} \rangle^2}{\langle P^{3P} \rangle^3} \qquad (23).$$

For Eq. (22) and (23) we assumed that the concentration within each measured cell during the consecutive two- and three-photon excitation measurements at 1100 nm, 80 MHz and 1650 nm, 3 MHz remains the same.

#### 1.2. Dependence of fluorescence signal and higher harmonics generation signals on the laser power

The Eq. (20) for the fluorescence signal upon non-resonant two-photon excitation in logarithmic representation is given by:

$$\lg\left(F_{2P}\right) = \lg\left(\frac{1}{2}\bar{\varphi}\cdot\sigma_{2P}\cdot\eta_{2P}\cdot C\cdot\frac{g_P{}^{2P}}{RR_{2P}\cdot\tau_p{}^{2P}}\frac{\pi\cdot n}{\lambda_{exc}}\right) + 2\lg\left(\langle P^{2P}\rangle\right) \quad (24).$$

In analogy, the second harmonics generation (SHG) signal in logarithmic representation is given by Eq. (25). The difference between Eq. (24) and (25) is due to the fact that the SHG signal depends on the real part of the complex second order susceptibility $\chi_2$, whereas the fluorescence signal upon a two-photon excitation depends on its imaginary part[7].

$$\lg\left(SHG\right) = \lg\left(\frac{1}{2}RE(\chi_2)\cdot\frac{g_P{}^{2P}}{RR_{2P}\cdot\tau_p{}^{2P}}\frac{\pi\cdot n}{\lambda_{exc}}\right) + 2\lg\left(\langle P^{2P}\rangle\right) \quad (25)$$

*RE($\chi_2$)* is the real part of the second order susceptibility.

In the same line, the fluorescence signal upon non-resonant three-photon excitation given by Eq(21) in logarithmic representation turns into:

$$\lg\left(F_{3P}\right) = \lg\left(\frac{1}{3}\bar{\varphi}\cdot\sigma_{3P}\cdot\eta_{3P}\cdot C\cdot\frac{g_P{}^{3P}}{RR_{3P}{}^2\cdot\left(\tau_p{}^{3P}\right)^2}\cdot 0.544\cdot\frac{\pi^2\cdot n\cdot NA^2}{\lambda_{exc}{}^3}\right) + 3\lg\left(\langle P\rangle\right) \quad (26).$$

The third harmonics generation (THG) signal in logarithmic representation is given by:

$$\lg\left(THG\right) = \lg\left(\frac{1}{3}RE(\chi_3)\cdot\frac{g_P{}^{3P}}{RR_{3P}{}^2\cdot\left(\tau_p{}^{3P}\right)^2}\cdot 0.544\cdot\frac{\pi^2\cdot n\cdot NA^2}{\lambda_{exc}{}^3}\right) + 3\lg\left(\langle P\rangle\right) \quad (27)$$

with *RE($\chi_3$)* the real part of the third-order susceptibility[7].

For a resonance-enhanced three-photon excitation (RE3P), we expect the fluorescence signal to be a linear combination of a two-photon excitation process given by Eq. (20) and a three-photon excitation process given by Eq. (21), with the pre-factors $a_{2P}$ and $a_{3P}$, respectively, the two pre-factors summing up to unity. In logarithmic representation, the fluorescence signal upon resonance-enhanced three-photon excitation is:

$$\lg\left(F_{RE3P}\right) = \lg\left[a_{2P}\cdot\left(\frac{1}{2}\bar{\varphi}\cdot\sigma_{2P}\cdot\eta_{2P}\cdot C\cdot\frac{g_P{}^{2P}}{RR_{2P}\cdot\tau_p{}^{2P}}\frac{\pi\cdot n}{\lambda_{exc}}\cdot\langle P\rangle^2\right) + a_{3P}\cdot\left(\frac{1}{3}\bar{\varphi}\cdot\sigma_{3P}\cdot\eta_{3P}\cdot C\cdot\frac{g_P{}^{3P}}{RR_{3P}{}^2\cdot\left(\tau_p{}^{3P}\right)^2}\cdot 0.544\cdot\frac{\pi^2\cdot n\cdot NA^2}{\lambda_{exc}{}^3}\cdot\langle P\rangle^3\right)\right] \quad (28).$$

With increasing laser excitation power, $a_{3P}$ increases, turning Eq. (28) into Eq. (26) for non-resonant three-photon excitation, dependent only on the cubic power of the average laser power.

#### 1.3. Laser power attenuation with tissue imaging depth (z)

As previously described, we can assume that the average laser power in a multi-photon setup exponentially decreases with tissue imaging depth z, mainly due to absorption and scattering of radiation:

$$\langle P\rangle(z) = \langle P\rangle(z=0)\cdot e^{-z/l_e} \quad (29)$$

with $\langle P\rangle(z=0)$ the average laser power at the surface of the specimen and $l_e$ the effective attenuation length.

When combining Eq. (20) and (29), the depth-dependent fluorescence signal upon two-photon excitation, in natural logarithmic representation, is given by:

$$\ln\left(F_{2P}(z)\right) = \ln\left(\frac{1}{2}\bar{\varphi}\cdot\sigma_{2P}\cdot\eta_{2P}\cdot C\cdot\frac{g_P{}^{2P}}{RR_{2P}\cdot\tau_p{}^{2P}}\frac{\pi\cdot n}{\lambda_{exc}}\right) + 2\ln(\langle P(z=0)\rangle) - \frac{2z}{l_e} \quad (30).$$

In analogy, the depth-dependent SHG signal is given by:

$$\ln(SHG) = \ln\left(\frac{1}{2}RE(\chi_2)\cdot\frac{g_P{}^{2P}}{RR_{2P}\cdot\tau_p{}^{2P}}\frac{\pi\cdot n}{\lambda_{exc}}\right) + 2\ln(\langle P(z=0)\rangle) - \frac{2z}{l_e} \quad (31).$$

The depth-dependent fluorescence signal upon three-photon excitation, also in natural logarithmic representation, is obtained when replacing Eq. (29) in Eq. (21):

$$\ln(F_{3P}) = \ln\left(\frac{1}{3}\bar{\varphi}\cdot\sigma_{3P}\cdot\eta_{3P}\cdot C\cdot\frac{g_P{}^{3P}}{RR_{3P}{}^2\cdot\left(\tau_p{}^{3P}\right)^2}\cdot 0.544\cdot\frac{\pi^2\cdot n\cdot NA^2}{\lambda_{exc}{}^3}\right) + 3\ln(\langle P(z=0)\rangle) - \frac{3z}{l_e} \quad (32).$$

For the THG signal, the equation reads:

$$\ln(THG) = \ln\left(\frac{1}{3}RE(\chi_3)\cdot\frac{g_P{}^{3P}}{RR_{3P}{}^2\cdot\left(\tau_p{}^{3P}\right)^2}\cdot 0.544\cdot\frac{\pi^2\cdot n\cdot NA^2}{\lambda_{exc}{}^3}\right) + 3\ln(\langle P(z=0)\rangle) - \frac{3z}{l_e} \quad (33).$$

As we used laser power adaptation with depth (z) in our measurements, the power at surface *P(z=0)* is also a function of z. Both fluorescence and higher-harmonics generation signals were normalized for the corresponding *P(z=0)* at the respective imaging depth.

#### 1.4. Fluorophore saturation limits the maximum imaging depth in tissues for a given pulsed excitation source

In order to determine the supported repetition rate for a non-resonant (multi-photon) excitation, in a certain imaging depth in tissue, for a given average laser power at the tissue surface (z = 0), we rely on the equation[7]:

$$RR_{3P} = \frac{\langle P(z=0)\rangle}{E_{focus}} \cdot e^{-z/l_e} \qquad (34),$$

with $E_{focus}$ the pulse energy at the focus.

If $E_{focus}$ exceeds the pulse energy $E_{sat}$, which leads to the saturation of the fluorophore, the fluorescence signal will not further increase. Experimentally, we avoid reaching the saturation regime, as shown by the double-logarithmic representations in Suppl. Fig. 2. In this way, we avoid a disproportionate increase of highly non-linear photobleaching, without increase of fluorescence signal.

For a three-photon excitation process[8], the saturation energy is given by: $E_{sat} = \frac{h \cdot c \cdot \lambda_{exc}}{\pi \cdot NA^2} \cdot \sqrt[3]{\frac{(\tau_p{}^{3P})^2}{g_P{}^{3P} \cdot \sigma_{3P} \cdot \eta_{3P}}}$. Hence, the maximum repetition rate is given by:

$$RR_{3P} = \frac{\langle P(z=0)\rangle}{\frac{h \cdot c \cdot \lambda_{exc}}{\pi \cdot NA^2} \cdot \sqrt[3]{\frac{(\tau_p{}^{3P})^2}{g_P{}^{3P} \cdot \sigma_{3P} \cdot \eta_{3P}}}} \cdot e^{-z/l_e} \qquad (35).$$

#### 1.5. Signal-to-noise ratio

The signal-to-noise ratio (SNR) in all cases, i.e. fluorescence upon non-resonant two-, non-resonant three-photon, or resonance enhanced three-photon excitation, as well as second and third harmonics generation, is defined as:

$$SNR = \frac{detected\ gray\ value - \langle background\rangle}{\sigma(background)} \qquad (36).$$

We thereby assume that the background (recorded in regions with no tissue structure) is Gaussian distributed, with $\langle background\rangle$ the mean background value and $\sigma(background)$ the width of the Gaussian function. $\sigma(background)$ represents the background noise.

### 2. Microscope setup for *in vivo* deep-tissue tibia imaging: design and characterization

We extended a state-of-the-art two-photon microscope to operate up to 1700 nm and allow for efficient three-photon excitation (Fig. 1). A detailed description of the microscope and its components can be found in *Methods*. We ensured that the selected optical elements of the microscope setup and their anti-reflex coatings are adequate for the extended infrared wavelength range, resulting into a total transmittance of the microscope system at 1650 nm of 28.3±0.4 %.

In order to compensate the slightly positive group-velocity dispersion (GVD) of our setup when using OPA lasers as excitation sources, we integrated either a two-prism pulse-compressor (for the wavelength-tunable OPA) or ZnSe windows[9] of variable thickness (for Ytterbia OPA). We found that a total thickness of 13.5 mm ZnSe provides the narrowest pulse width under the objective lens. The optimum ZnSe thickness and the performance of the pulse compressor were assessed by measuring the pulse width by second-order interferometric autocorrelation under the objective lens. While not counteracting the GVD compensation at the level of prism-based pulse compressors, the loss of laser power through ZnSe material is lower compared to state-of-the-art prism pulse compressors, by a reduced number of optical elements in the beam path. This ensured a higher average power, i.e. pulse-energy, under the objective lens. Additionally, we also avoided higher-order dispersion, i.e. possible distortions of the temporal pulse shape.

Further, we tested the suitability of two objective lenses, specifically designed for infrared multi-photon imaging up to 1600 nm (Olympus, XLPLN25XWMP2 and Nikon, CFI75 Apo25XCW1300). Therefore, we measured the pulse width of the tunable OPA laser source under each objective lens. We found similar values for both lenses, increasing from 55 fs up to 90 fs between 1300 and 1700 nm (Suppl. Fig. 1), using a two-prism pulse compressor to counteract the GVD of the microscope setup. However, the higher transmission at longer wavelengths (between 1600 and 1700 nm) was decisive for the use of the XLPLN25XWMP2 objective in all further experiments. The back aperture, for both objective lenses, was overfilled by a factor of 1.5 to achieve best spatial resolution.

### 2.1. Spatial resolution of the microscope setup

High-resolution 3D imaging of 100 nm nanospheres ($\lambda_{em}$ 605 nm) embedded in agarose (imaged volume 442x442x1031 $\mu m^3$, 1036x1036x1031 voxel) upon excitation at 1650 nm (3.09 MHz) indicate homogeneous illumination of the entire field of view (FOV), over more than 1 mm depth (Suppl. Fig. 1). Similar lateral (xy) and axial (z) PSF line profiles, in different regions within the full FOV (Suppl. Fig. 1) and at different imaging depths (Suppl. Fig. 1), indicate negligible or no wave front distortions or loss of spatial resolution, caused by the optical elements.

### 2.2. Characterization of multi-photon processes leading to fluorescence and higher harmonics generation

A prerequisite to characterize the performance of the distinct laser systems as excitation sources for *in vivo* deep-tissue tibia imaging is to assess the type of excitation, two- or three-photon excitation, and excitation capacity, i.e. absorption cross-section, for the fluorescent proteins expressed in the reporter mice used for imaging.

We verified the type of non-linear excitation process leading to fluorescence for GFP, tdRFP and tdTomato (expressed in Cdh5:tdTomato x Histone:GFP (Cdh5:tdTom) , CD19:tdRFP or Prx1:tdRFP mice used in this study), in both infrared spectral windows adequate for deep-tissue imaging, i.e. at 1330 nm and 1650 nm. We calculated the slope of the fluorescence signal dependence on the average laser power, in double logarithmic representation, to differentiate between two- and three-photon excitation processes, based on Eq. (24), (26), (28). At 1330 nm (Suppl. Fig. 2), we found for GFP fluorescence a slope of ≈3, complying with a non-resonant three-photon excitation, whereas for tdTomato fluorescence the dependence on the average laser power was not linear, with a slope <3 for low laser powers and ≈3 for high laser powers. This indicates a resonance-enhanced three-photon excitation, i.e. a two-step 2+1 photon excitation, in agreement with previously published work[3]. At 1650 nm (Suppl. Fig. 2), we found slopes of ≈3 for both tdTomato and tdRFP fluorescence, indicating non-resonant three-photon excitation processes. As expected, we found a slope of ≈2 for the SHG signal (in the collagen fibers of bone cortex) and of ≈3 for the THG signal (in lacunae and canaliculi of the bone cortex), at 1650 nm (Suppl. Fig. 2).

#### 2.3. Active three-photon excitation cross-sections of tdRFP and tdTomato at 1650 nm

The active three-photon absorption cross-section $\eta_3 \cdot \sigma_3$ is a key molecular parameter for estimating the excitation efficacy and, thus, the three-photon imaging performance of the respective fluorophore in tissue. Based on the ratio of the fluorescence signals generated upon two-photon excitation at 1100 nm and three-photon excitation at 1650 nm, respectively (Eq. (22)), we calculated the active three-photon absorption cross-section $\eta_3 \cdot \sigma_3$ at 1650 nm for both tdRFP and tdTomato, using published values of the active two-photon absorption cross-section $\eta_2 \cdot \sigma_2$. We estimated for $\eta_3 \cdot \sigma_3$ of tdRFP expressed in B lymphocytes (from a CD19:tdRFP mouse) a value of $15.85 \pm 0.2 \cdot 10^{-83}$ $cm^6 \cdot s^2$ at 1650 nm, 3.09 MHz, 106 fs pulse width. Two-photon excitation was performed in the same cells at 1100 nm, 80 MHz, 166 fs pulse width, with $\eta_2 \cdot \sigma_2$ for tdRFP $20.2 \cdot 10^{-50}$ $cm^4 \cdot s$ (20.2 GM)[10]. For tdTomato in endothelial cells (Cdh5:tdTom) we estimated for $\eta_3 \cdot \sigma_3$ a value of $21.64 \pm 1.35 \cdot 10^{-83}$ $cm^6 \cdot s^2$ at 1650 nm, 3.09 MHz, 106 fs pulse width. We performed two-photon excitation in the same cells at 1100 nm, 80 MHz, 166 fs pulse width, with $\eta_2 \cdot \sigma_2$ for tdTomato considered to be $80 \cdot 10^{-50}$ $cm^4 \cdot s$ (80

GM)[11]. The calculated $\eta_3 \cdot \sigma_3$ value for tdTomato, while estimated in cellular environment, has the same order of magnitude as previously reported $\sigma_3$[3].

### 3. Parameter optimization of 3PM setup for *in vivo* deep-marrow imaging in tibia

#### 3.1. Imaging through thin bone cortex in explanted tibia: performance of high pulse energy 1650 nm 3PM vs. low pulse energy 1100 nm 2PM

In order to assess the imaging performance upon three-photon compared to two-photon excitation in the tibia marrow, we analyzed the depth-dependent SNR (ddSNR) of tdTomato fluorescence in the thinned tibia of a Cdh5:tdTom mouse (12 weeks, cortical thickness after thinning ≈70 µm) at 1650 nm (3 MHz) and at 1100 nm (80 MHz), respectively (Suppl. Fig. 4). The ddSNR measured upon 1100 nm excitation declines to a value of 1 in ≈160 µm depth in the tibia marrow, measured from the interface between bone cortex and bone marrow. In contrast, upon three-photon excitation at 1650 nm (3 MHz, Ytterbia OPA) the ddSNR reaches the value 1 in ≈300 µm depth in the tibia marrow, being by a factor of ≈1.9 larger (Suppl. Fig. 4). For comparability reasons, we adjusted the average laser power of Ytterbia OPA and OPO to achieve the same SNR of tdTomato fluorescence using both excitation schemes at the interface between bone cortex and bone marrow. Further, we performed imaging without z-adaptation of power, at the same pixel dwell time (1.98 µs, 2x frame acquisition). We could not compare the ddSNR using the two excitation schemes in the intact tibia of adult mice, because imaging through the thick intact bone cortex is not possible at low pulse energy, as shown above.

Further, we succeeded to image tdTomato fluorescence in blood vessels, in the marrow tissue underneath the intact bone cortex of the freshly explanted tibia of a 106 weeks old Cdh5:tdTom mouse at both 1650 nm (3 MHz) and at 1100 nm (80 MHz) (Suppl. Fig. 4), as the cortical bone of frail female mice is locally thinner than in adult mice[12]. These observation was confirmed in the explanted tibia bones of two further 127 weeks old Cdh5:tdTom mice.

#### 3.2. Impact of pulse energy on imaging depth in intact tibia *in vivo*

In order to verify that high pulse-energy is central for successful imaging through intact tibia cortex, we performed imaging in the distal area of the tibia, in a Cdh5:tdTom mouse, at 1330 nm, 2 MHz (tunable OPA) and at 1330 nm, 80 MHz (OPO), respectively. The average power of both laser sources was adjusted to 30 mW (without adaptation in z-direction) and the pixel dwell time was kept the same, namely 2.46 µs (2x frame acquisition) for both excitation schemes. Only the pulse energy differed, amounting to 15 nJ at 1330 nm, 2 MHz OPA illumination and 0.375 nJ at 1330 nm, 80 MHz OPO illumination. Imaging of SHG, THG, tdTomato and GFP fluorescence signals in the tibia (cortex and marrow) was achieved with the high, but not with the low pulse energy (Suppl. Fig. 4). At low pulse energy (OPO), only THG and SHG signals of the upper tissue layers in the bone cortex could be detected. At the higher pulse energy (tunable OPA), next to the SHG and THG signals originating from bone tissue and tdTomato and GFP fluorescence from the membranes and nuclei of endothelial cells in the bone marrow, we detected a strong SHG signal in connective tissue and muscles adjacent to the tibia.

#### 3.3. Physical parameters limiting maximum imaging depth in tibia bones

The maximum laser repetition rate in tissue may be limited by the saturation energy of the fluorophore, in our case tdTomato or tdRFP, and by the maximum average power at the surface of the specimen[8,13]. Using the parameters of the Ytterbia OPA, we calculated the maximum repetition rate in tibia following Eq. (35), relying on the saturation energy values of 4.03 nJ for tdTomato and of 4.47 nJ for tdRFP, which were calculated from the previously determined active three-photon absorption cross-section $\eta_3 \cdot \sigma_3$ values of the two fluorescent proteins. In 500 µm depth in intact tibia, hereby assuming 150 µm cortical thickness and 350 µm imaging depth in the bone marrow, the maximum repetition rate supported by an average power of 500 mW at the surface of the bone cortex is 7.83 MHz for tdRFP and 18.74 MHz for tdTomato. Thus, the highest repetition rate of 3.98 MHz, available in our system, is still supported by the maximum average power under the objective lens, i.e. 512 mW, for both tdRFP and tdTomato. Conversely, at 3.98 MHz and 512 mW, the maximum imaging depth in bone marrow, through 150 µm bone cortex, is 564 µm for tdRFP and 818 µm for tdTomato.

Thus, we conclude that the maximum available average power of our laser, i.e. Ytterbia OPA, rather than the repetition rate leading to saturation of tdRFP and tdTomato three-photon excitation retains the potential to limit the maximum imaging depth in mouse tibia. However, to prevent tissue photodamage and fluorophore photobleaching, we did not exceed 120 mW (at 3 MHz) in time-lapse tibia imaging experiments *in vivo*, and, by doing so, we avoided to saturate the fluorescent proteins.

#### 3.4. Photodamage analysis of spleen tissue by *in vivo* 3PM at 1650 nm

We performed photodamage analysis upon repetitive *in vivo* irradiation at 1650 nm not only in the organ of interest, the tibia, but also in spleen, as we expect this lymphoid organ to be more prone to photodamage than the tibia bone marrow, due to lack of protection by calcified bone. Additionally, controlled bone marrow damage by excessive laser irradiation at 850 nm, 80 MHz in intact tibia of adult mice was not possible, due to scattering and absorption of 850 nm radiation in calcified bone.

Using immunofluorescence analysis, we investigated possible tissue damage induced by repeated 1650 nm irradiation in spleen, as compared to non-irradiated samples and samples damaged by excessive laser irradiation at 850 nm (80 MHz), >300 mW (Suppl. Fig. 6). The spleens were continuously exposed over 10 min, under *in vivo* conditions, to 1650 nm radiation, average power 70 mW, at 2.06 MHz repetition rate, corresponding to a pulse energy of 34 nJ. Any increase in the number of neutrophils (Ly6G), or macrophages (CD68), which are known to respond in a rapid manner to (laser) damage, by migrating into the respective regions[14] was observed (Suppl. Fig. 6). Similarly, there was no evidence of increased apoptosis analyzed by TUNEL staining, or of cells expressing heat shock protein 70 (HSP70)[15,16] at the tissue sites irradiated at 1650 nm, compared to non-irradiated controls. In contrast, intentionally photodamaged spleen tissue showed an increase in the number of neutrophils, $TUNEL^+$ and $HSP70^+$ cells (Suppl. Fig. 6).

#### 3.5. tdRFP photobleaching during *in vivo* 3PM tibia imaging at 1650 nm

Next to tissue photodamage, fluorophore photobleaching, expected to scale highly non-linearly with the laser power[10,17], may limit the suitability of 1650 nm excitation for time-lapse *in vivo* applications. To investigate fluorophore photobleaching upon excitation at 1650 nm, we performed *in vivo* time-lapse imaging of tibia marrow in CD19:tdRFP and Prx1:tdRFP mice. In CD19:tdRFP fate mapping mice, tdRFP is expressed in B lineage cells, in Prx1:tdRFP fate mapping mice, tdRFP is expressed in mesenchymal stromal cells (MSC), and their progeny, such as osteoblasts, osteocytes or adipocytes[18].

To evaluate the tdRFP photobleaching induced upon three-photon excitation at 1650 nm, we directly compared it to photobleaching induced by well characterized two-photon excitation at 1100 nm. Therefore, *CD19:tdRFP* mice were imaged first upon two-photon excitation at 1100 nm, 80 MHz (OPO), followed by three-photon excitation at 1650 nm, 3 MHz (Ytterbia OPA), shown in Suppl. Fig. 7. Time-lapse imaging was performed every 30 s, over 60 minutes in 2PM regime, followed by 60 minutes in 3PM regime. By measuring the tdRFP fluorescence over time, we determined a photobleaching rate of $k_{photobl} = 2.4\cdot10^{-3}$ $min^{-1}$ for 2PM (1100 nm, 80 MHz, 0.31 nJ, 25 mW) and $k_{photobl} = 1.3\cdot10^{-3}$ $min^{-1}$ for 3PM (1650 nm, 3 MHz, 12.33 nJ, 37 mW). Thus, we concluded that 3PM induces similarly low tdRFP photobleaching as 2PM.

Fast time-lapse (every 0.5 s, over 3 minutes and every 1 s, over 3 minutes) imaging of tdRFP fluorescence the tibia marrow of Prx1:tdRFP mice, upon excitation at 1650 nm, 3 MHz repetition rate, 50 mW average power and 16 nJ pulse energy was performed to determine the photobleaching rate at fast image acquisition speed, over short imaging time windows (Suppl. Fig. 7). Additionally, we determined the tdRFP photobleaching rate at slow image acquisition speed and over long time periods, i.e. every 30 s over 60 minutes or every 120 s over 120 min, by performing time-lapse *in vivo* imaging in the tibia CD19:tdRFP mice at 1650 nm, 3MHz, at the average laser power and pulse energy indicated in Suppl. Fig. 7. We found that the tdRFP photobleaching rate increases when the time step was reduced, and the image acquisition speed increased. We determined $k_{photobl} = 9.85 \cdot 10^{-2}$ $min^{-1}$ for a time step of 0.5 s, $k_{photobl} = 6.92 \cdot 10^{-2}$ $min^{-1}$ for 1 s, $k_{photobl} = 1.3 \cdot 10^{-3}$ $min^{-1}$ for 30 s, and $k_{photobl} = 0.56 \cdot 10^{-3}$ $min^{-1}$ for 120 s.

**Supplemental Figures**

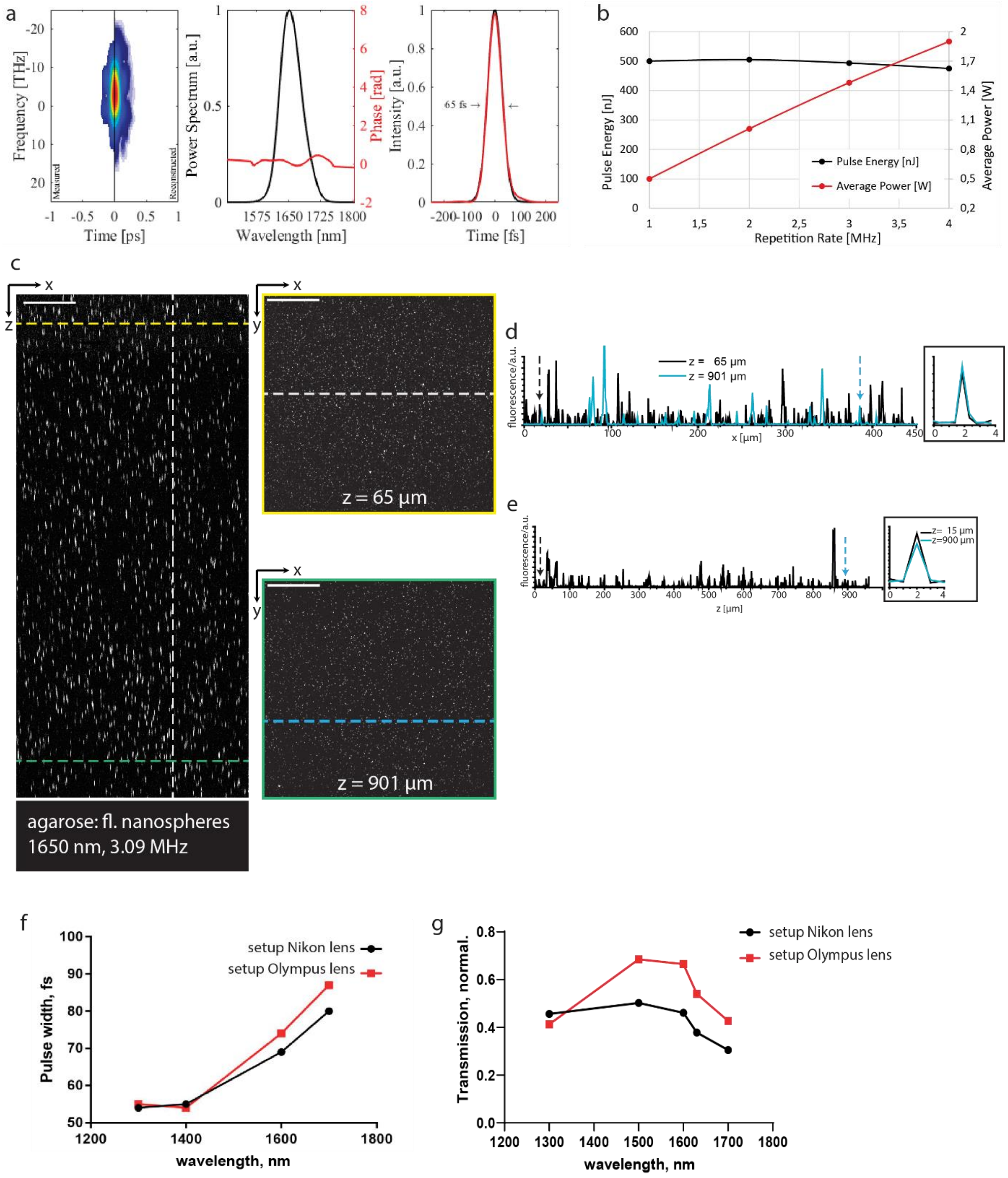


**Supplemental Figure 1: Characterization of the optical parametric amplifier with tunable repetition rate and of the multi-photon microscope setup used for three-photon imaging of intact murine tibia. a.** The second harmonic frequency resolved optical gating characterization of the Ytterbia optical parametric amplifier (OPA) pulse. (i) The Phantom-FROG trace of the 1650 nm pulse, with the measured trace on the left and the reconstructed trace on the right side of the graph. (ii) The reconstructed power spectrum (black) and phase (red). (iii) The reconstructed intensity profile (red), scaled to the normalized transform limited intensity profile (black) of the power spectrum. **b**. The pulse energy (black) and average power (red) scaling of the Ytterbia OPA as a function the repetition rate. **c**. xz (442x1001 µm², 1036x1001 pixel) and corresponding xy (442x442 µm², 1036x1036 pixel) fluorescence images of 100 nm fluorescent nanospheres embedded in agarose (emission at 605 nm), upon excitation at 1650 nm OPA radiation, at 3.09 MHz repetition rate, 0.8 nJ pulse energy. The horizontal yellow line in the xz fluorescence image shows the z-position (65 µm) of the yellow-framed xy image, the horizontal green line, the z-position (901 µm) of the green-framed xy image. The vertical white line in the xz-image shows the x-position of the fluorescence signal profile in e. The horizontal white line in the yellow-framed xy image shows the black profile in d, the cyan line in the green-frame xy image, the cyan profile also in d. **d**. Line profiles along x, as indicated in c, showing similar signal strength along the x-axis. The black arrow indicates the x-profile of a single sphere in 65 µm depth, the cyan arrow the x-profile of a single sphere in 901 µm depth. These two profiles are displayed in the inset of d, showing no loss of lateral resolution with depth. **e**. Line profile along z, as indicated in c, showing also axially similar signal strength. The black and cyan arrows indicate the z-profiles of single nanospheres, in 15 and 901 µm depth, respectively, which are displayed in the inset of e, showing also no loss of axial resolution with depth. **f**. Dependence of pulse width of the tunable OPA laser on the wavelength, measured under two different objective lenses (Olympus, XLPLN25XWMP2, 25x, NA 1.05 and Nikon, CFI75 Apo 25XC W 1300, 25x, NA 1.1), which were designed for high transmission in the infrared range. **g**. Transmission of the setup using the same two objective lenses, in the wavelength range 1300 nm – 1700 nm. Scale bar = 100 µm in all images.

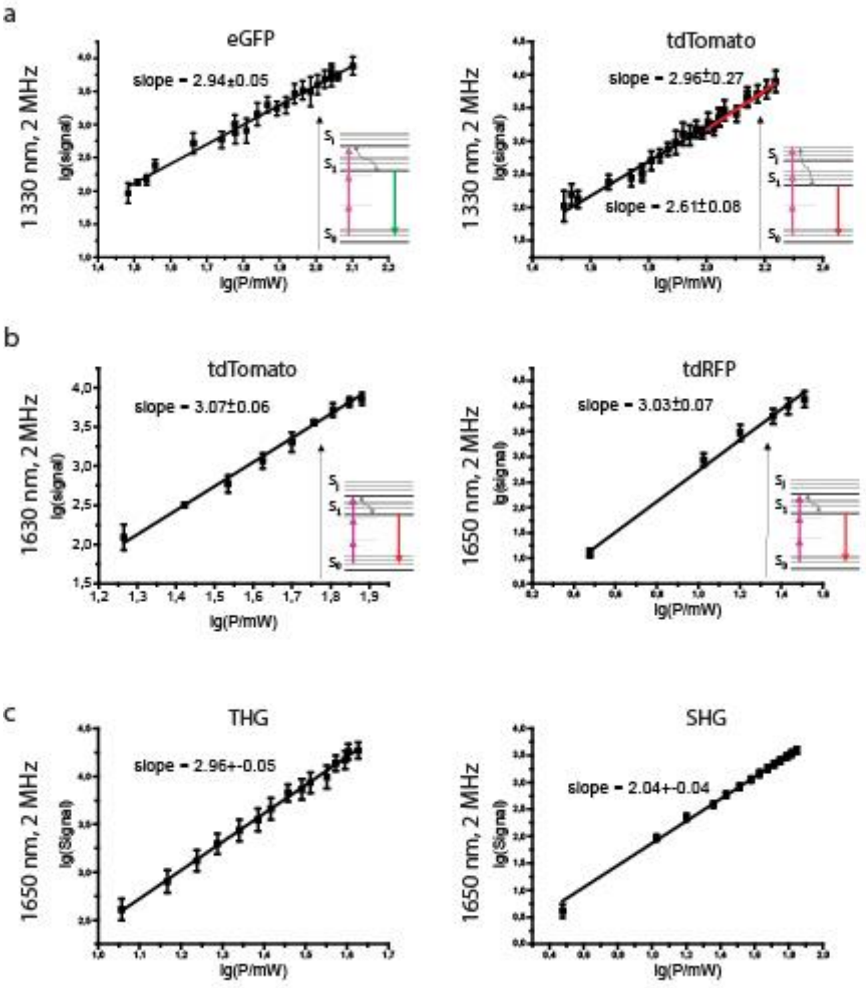


**Supplemental Figure 2: Wavelength dependence of the optically non-linear processes leading to fluorescence or higher harmonics generation. a**. Dependence of GFP and tdTomato fluorescence signal, respectively, on the average power of the OPA laser tuned at 1330 nm, in double-logarithmic representation. The slope of the linear approximation indicates the type of excitation, preceding spontaneous emission: a slope of 3 hints to a non-resonant three-photon excitation, a slope below 3, approaching 3 only at high powers, hints towards a resonance-enhanced three-photon excitation. Schematics of the energetic molecular levels and transitions are displayed as insets in graphs. **b**. As in a, dependence of tdTomato and tdRFP fluorescence signals on the laser average power at 1630 nm or at 1650 nm. In both cases, the slope of 3 indicates a non-resonant three-photon excitation, as shown also by the schematics of the energy diagram, as graph insets. **c**. Dependence of second harmonics generation (SHG) and third harmonics generation (THG) on the average laser power, at 1650 nm, 2.06 MHz (Ytterbia OPA).

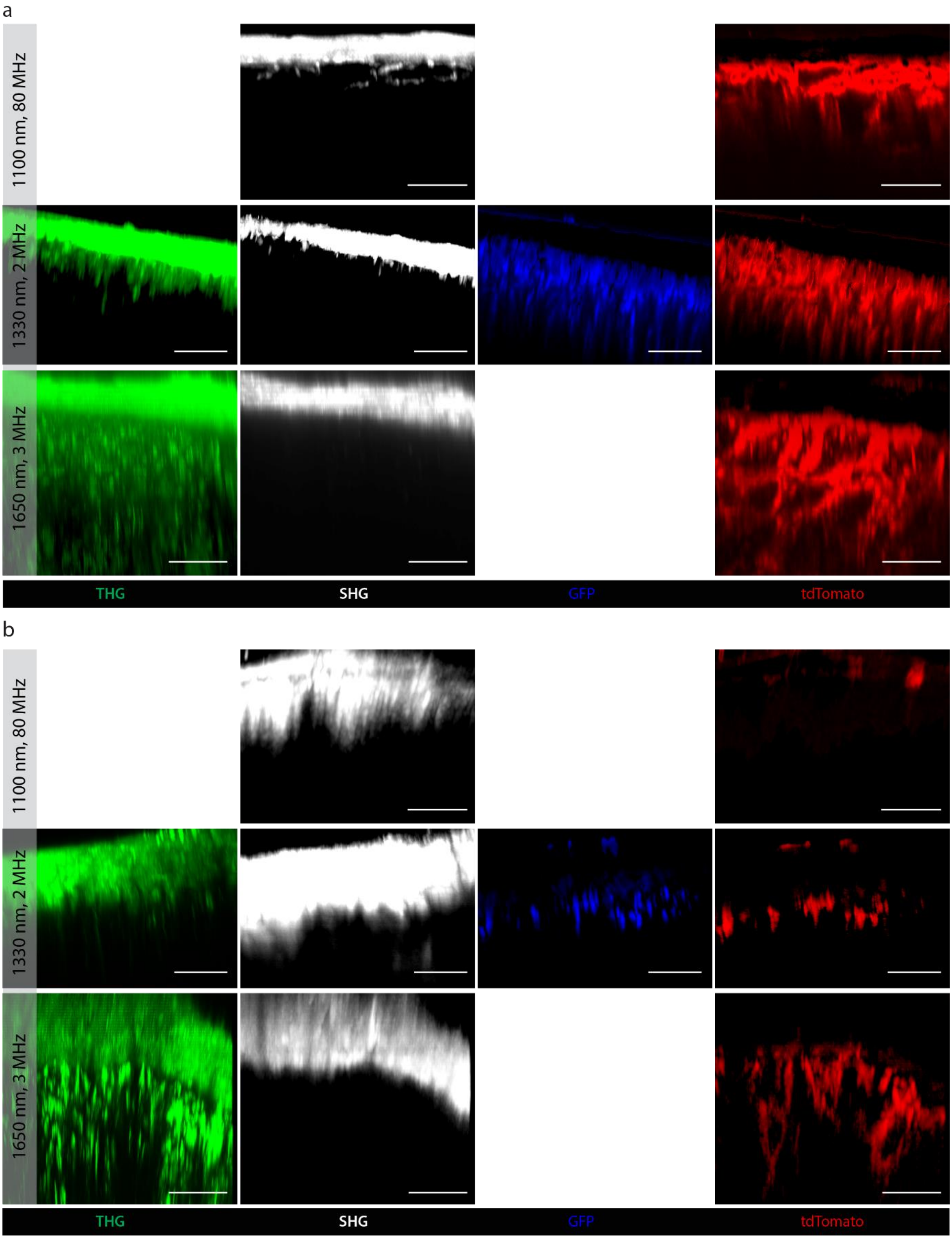


**Supplemental Figure 3: Single channel images in the tibia of adult mice, either with mechanically thinned or intact bone cortex, corresponding to the merged images in Figure 2. a.** Single channel z projections of 3D images acquired by 3PM at 1650 nm, 3 MHz or at 1330 nm, 2 MHz and by 2PM at 1100 nm, 80 MHz in the tibia of Cdh5:tdTomato x Histone:GFP mice, with mechanically thinned cortical bone. SHG signal is shown gray, THG signal in green, tdTomato fluorescence in the endothelial cell membranes in red and GFP fluorescence in the endothelial cell nuclei in blue. **b**. Single channel z projections of 3D images acquired by 3PM at 1650 nm, 3 MHz or at 1330 nm, 2 MHz and by 2PM at 1100 nm, 80 MHz in the intact tibia of Cdh5:tdTomato x Histone:GFP mice. Color coding as in **a**. Scale bar =100 µm.

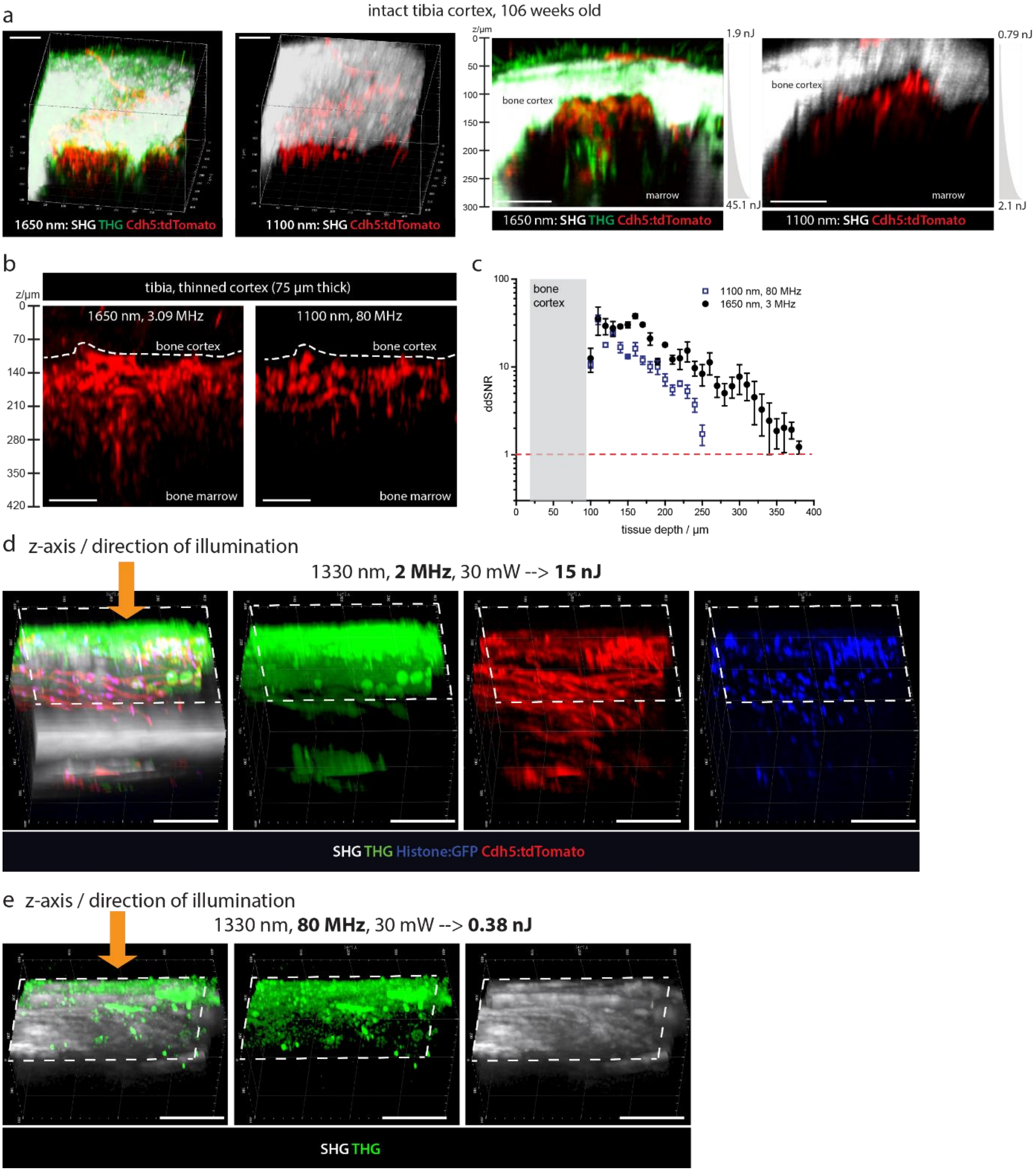


**Supplemental Figure 4: Both high-pulse energy and long wavelength excitation are required to maximize imaging depth in murine tibia.** **a**. 3D reconstructions (left side) and xz projections over 50 µm along the y axis (right side) of SHG (gray), THG (green) and tdTomato fluorescence (red) in blood vessels of a Cdh5:tdTomato x Histone:GFP mouse (106 weeks old, with locally thin bone cortex), upon excitation at 1650 nm, 3.09 MHz, and at 1100 nm, 80 MHz, respectively. For both excitation schemes exponential z-adaptation of laser power was used as indicated. **b**. xz projections over 50 µm in y-direction (400x400x420 µm³, 518x518x160 voxel) of tdTomato fluorescence in blood vessels in the tibia of a Cdh5:tdTomato x Histone:GFP mouse, with mechanically thinned cortex (75 µm thick), upon excitation at 1650 nm, 3.09 MHz and 1100 nm, 80 MHz, without z-adaptation of power. **c**. Depth dependence of SNR values calculated for the tdTomato fluorescence, in images shown in b. **d**. 3D rendered images (400x400x300 µm³, 518x518x150 voxel) of SHG (gray), THG (blue), GFP (green) and tdTomato (red) fluorescence in the intact tibia of a Cdh5:tdTomato x Histone:GFP mouse (12 weeks old), acquired at 1330 nm, 2 MHz (tunable OPA) or at 1330 nm, 80 MHz (OPO) in **e**. Both excitation sources (OPA and OPO) were adjusted at 30 mW, corresponding to pulse energies of 15 nJ for the OPA and 0.38 nJ for the OPO. The orange arrows in both d and e indicate the direction of illumination (the optical axis of the microscope). Scale bar = 100 µm for all images.

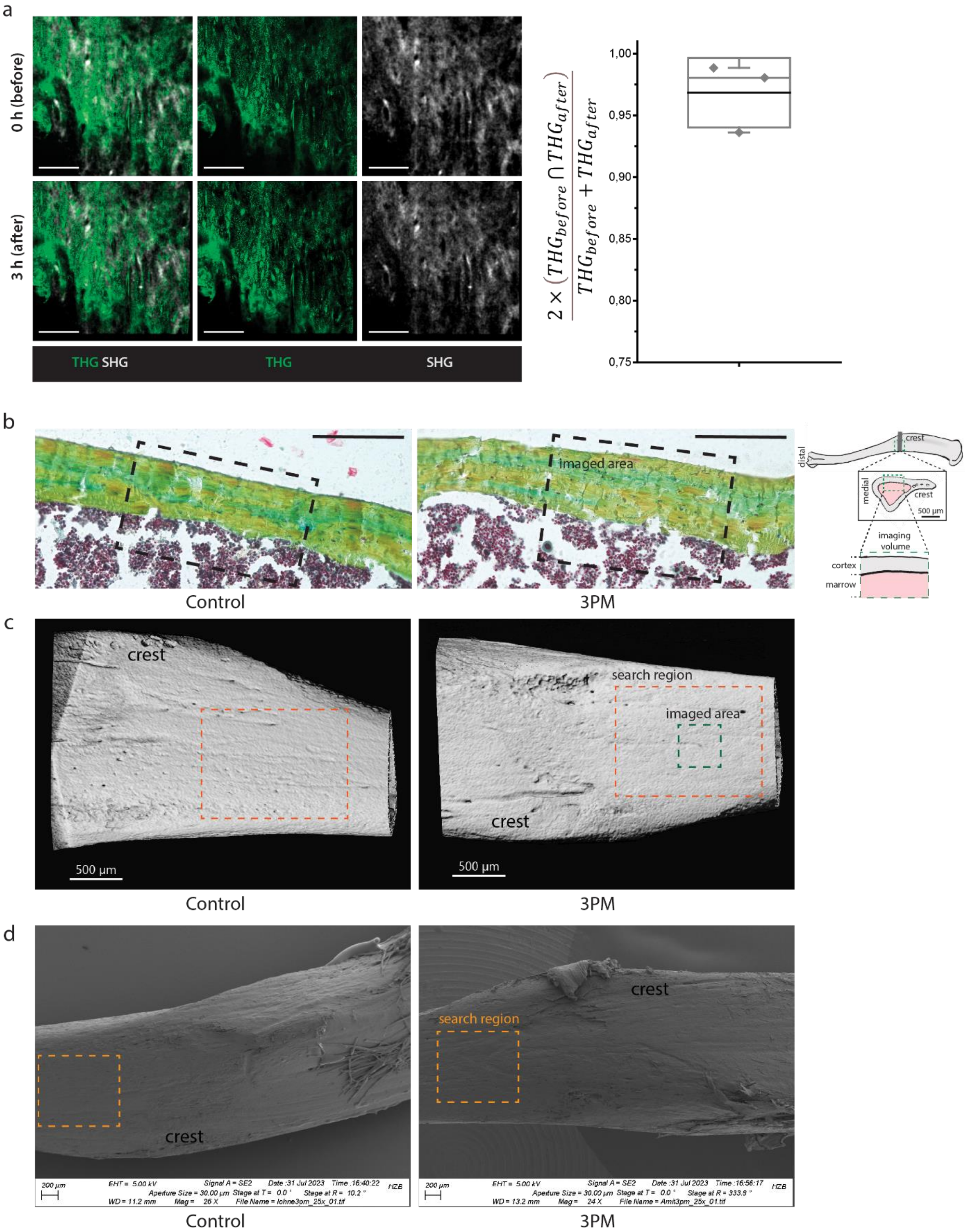


**Supplemental Figure 5: Analysis of the bone surface integrity after time-lapse intravital tibia imaging upon excitation with high pulse energy at 1650 nm. a.** Representative maximum intensity projections (30 µm thick) of THG and SHG signal at the tibia surface before and after intravital imaging at 1650 nm (3 MHz, 80 mW). The intravital imaging was performed in a time-lapse manner over 3 hours. Scale bar = 100 µm. The difference between the THG signal before and after intravital time-lapse imaging over 3 hours calculated as $2\left(THG_{before} \cap THG_{after}\right)/\left(THG_{before} + THG_{after}\right)$ is displayed for 3 CD19:tdRFP mice. **b**. Representative Movat's pentachrom staining of the bone cortex of a murine tibia exposed to 1650 nm, 3 MHz (80 mW) over 3 hours, during intravital time-lapse imaging. A non-exposed tibia was used as control. Scale bar = 200 µm. **c**. Representative NanoCT 3D reconstructions of a tibia exposed for 2 hours to 1650 nm laser radiation (3 MHz, 110 mW, 35.6 nJ) during intravital imaging in comparison to a non-exposed tibia. Search region (orange dashed rectangle) is region, from which the area to be imaged was chosen. Dark green dashed square is representative for the imaged area. Scale bar = 500 µm. **d**. Representative SEM images of the surface of a tibia exposed for 2 hours to 1650 nm laser radiation (3 MHz, 115 mW, 37.2 nJ) during intravital imaging (3PM) in comparison to a non-exposed tibia (Control). Similar to **c**, the orange dashed rectangle represents the search region. Scale bar = 200 µm.

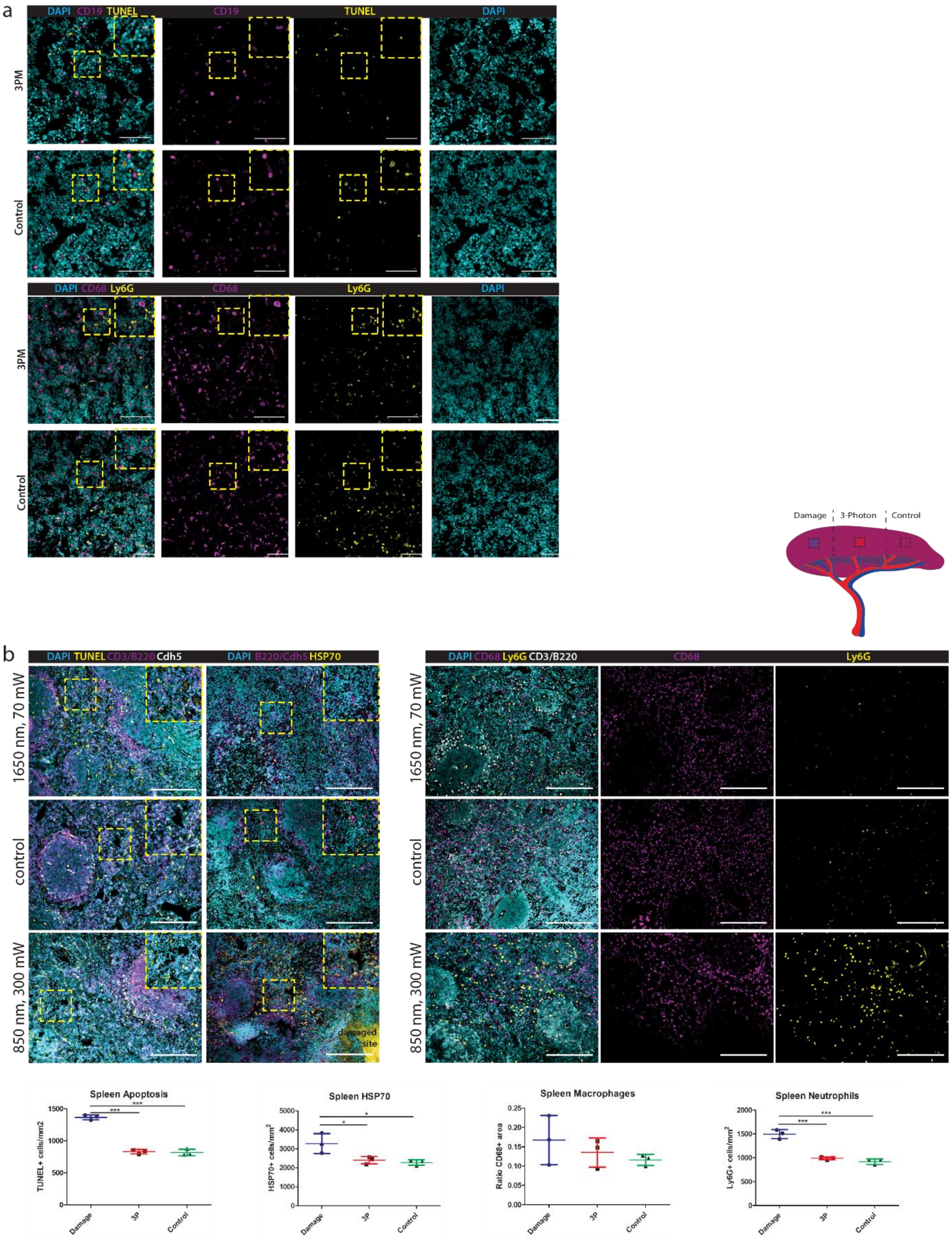


**Supplemental Figure 6: Immunofluorescence analysis of tissue photodamage after time-lapse imaging upon excitation with high pulse energy at 1650 nm in tibia marrow and spleen. a.** Representative immunofluorescence overlays of TUNEL (apoptosis, yellow), nuclear (DAPI, cyan) and CD19 (B lineage cells, magenta) and overlays of macrophages (CD68, magenta), neutrophil granulocytes (Ly6G, yellow), and nuclear (DAPI, cyan) staining, in tibia marrow tissue irradiated at high pulse energy 1650 nm (3.09 MHz, 120 mW), and not irradiated (control). Scale bar = 200 µm. **b.** Representative immunofluorescence overlays of TUNEL (apoptosis, yellow), nuclear (DAPI, cyan), CD3/B220 (white pulp, magenta) and Cdh5 (vessels, white) and overlays of HSP70 (yellow), nuclear (DAPI, cyan) and B220 (white pulp, magenta)/Cdh5(vessels, magenta), in spleen tissue irradiated at high pulse energy 1650 nm (2.06 MHz, 70 mW), excessively irradiated at 850 nm (80 MHz, 300 mW), and not irradiated. Representative immunofluorescence overlays of macrophages (CD68, magenta), neutrophil granulocytes (Ly6G, yellow), CD3/B220 (white pulp, white) and nuclear (DAPI, cyan) staining, in spleen tissue irradiated at high pulse energy 1650 nm (70 mW), excessively irradiated at 850 nm (300 mW), and not irradiated. Scale bar = 200 µm. Levels of apoptosis, heating damage (HSP70), numbers of macrophages and neutrophil granulocytes, in spleen tissue irradiated at 1650 nm, 3 MHz and measured by immunofluorescence histology (n = 3 mice).

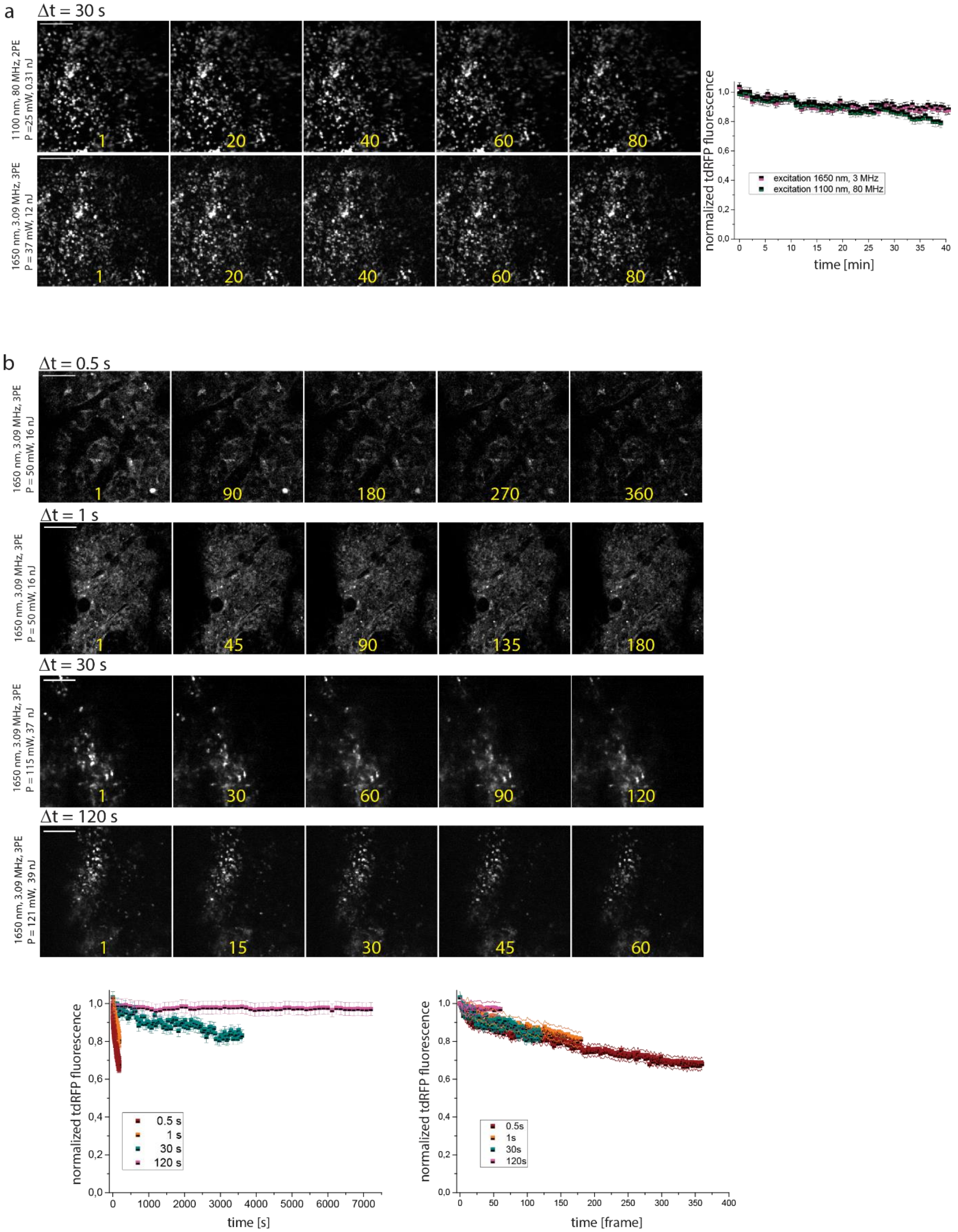


**Supplemental Figure 7: Photobleaching analysis during intravital time-lapse tibia imaging upon excitation with high-pulse energy 1650 nm radiation. a**. Intravital time-series of selected tdRFP fluorescence images in the tibia of a CD19:tdRFP mouse upon excitation at 1100 nm, 80 MHz (low pulse energy) and at 1650 nm, 3.09 MHz (high pulse energy). Scale bar 100 µm. Graphs represent the decay of the average tdRFP fluorescence signal during intravital imaging every 30 s, over 40 min, upon excitation at 1100 nm, 80 MHz and at 1650 nm, 3 MHz, respectively, at the indicated average powers and pulse energies. **b**. Intravital time-series of selected tdRFP fluorescence images in the tibia of Prx1:tdRFP (time step 0.5 s and 1 s) or CD19:tdRFP (time step 30 s and 120 s) mice upon excitation at 1650 nm, 3 MHz (high pulse energy). Scale bar 100 µm. Graphs represent the decay of the average tdRFP fluorescence signal during intravital tibia imaging for various time steps, i.e. 0.5 s, 1 s, 30 s, and 120 s, respectively.

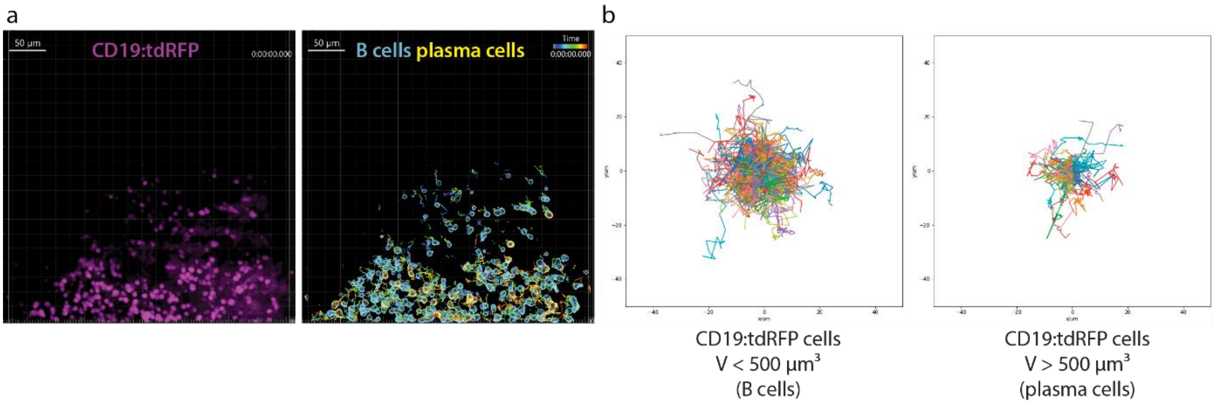


**Supplemental Figure 8: Intravital imaging of bone marrow B lineage cells in the intact tibia of a CD19:tdRFP mouse upon excitation at 1650 nm, 4 MHz. a.** Representative 3D reconstructions of tdRFP fluorescence signal (magenta) of B lineage cells (400x400x60 µm³, 518x518x11 voxels) and segmented cells, differentiating between B cells (volume < 500 µm³, cyan outlines) and plasma cells (volume > 500 µm³, yellow outlines). Scale bar = 50 µm. Imaging time window = 2 hours, time step = 120 s. **b**. Corresponding rose plots displaying the trajectories of B cells and plasma cells, respectively.

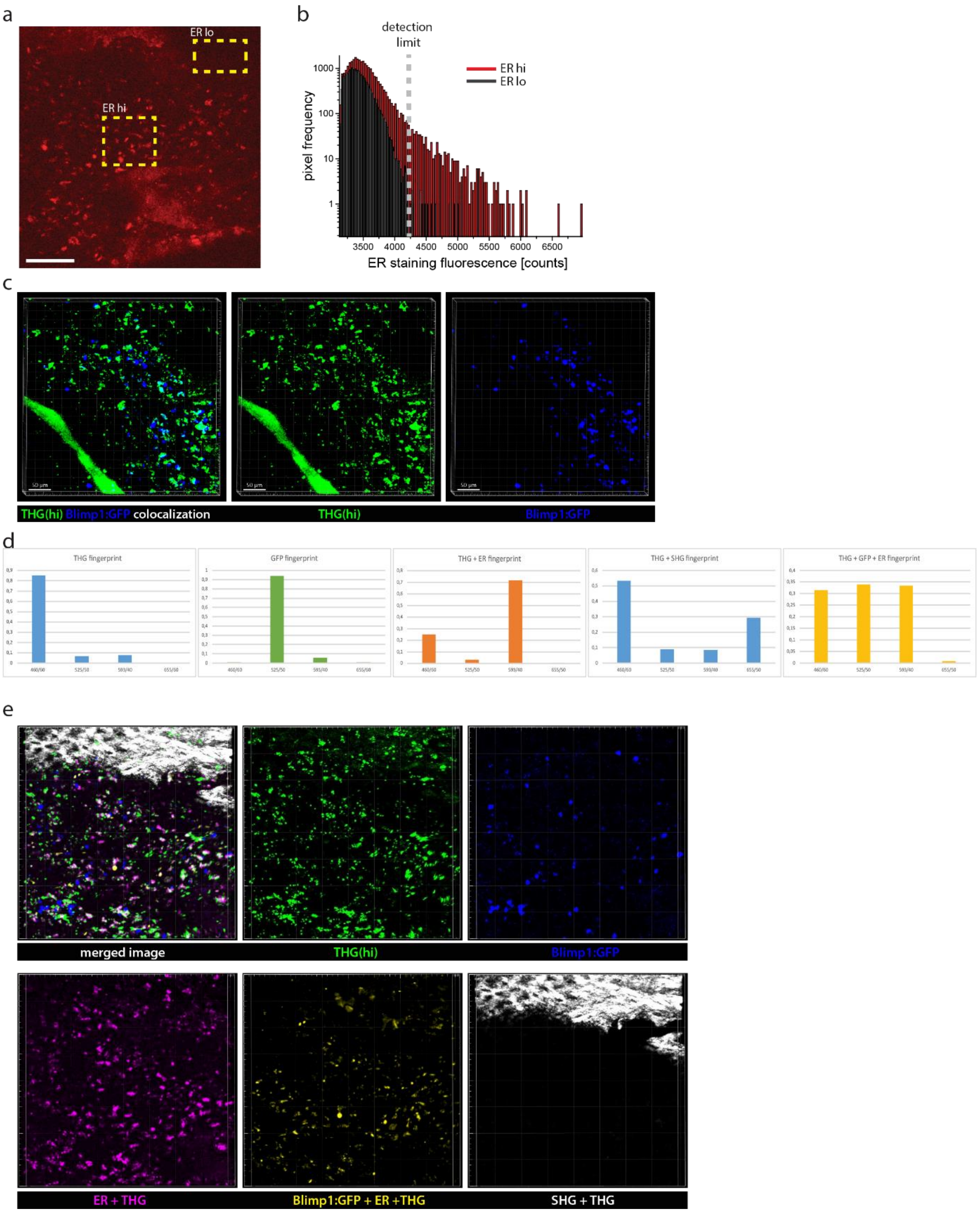


**Supplemental Figure 9: Analysis of endoplasmic reticulum (ER) staining in the tibia marrow. a.** Representative fluorescence image of ER staining in the bone marrow of a murine tibia. Scale bar = 100 µm. The yellow rectangles indicate a background region and a region with cells positively stained for ER. **b**. Histogram distributions of background and ER staining fluorescence signal, respectively. The upper limit of the background is defined as the limit for positive (abundant) ER staining, i.e. ER detection limit. **c**. 3D image (442x442x100 µm$^3$, 1010x1010x51 voxel) of THG (green) and GFP (blue) in the bone marrow of an explanted

Blimp1:GFP mouse tibia (3PM at 1330 nm). Scale bar = 50 µm. **d**. *In situ* finger prints for SIMI spectral unmixing in the tibia marrow of *Blimp1:GFP* mice, detected upon excitation at 1330 nm in the channels 460/60, 525/50, 593/40 and 655/50. We could identify pixels displaying only THG signal, THG and SHG signals, only GFP signal, THG and ER tracker signals and GFP, ER tracker and THG signals. **e**. Merged 3D image (400x400x60 µm$^3$, 518x518x31 voxel, upper left) and spectrally (using SIMI) unmixed tissue components in an explanted Blimp1:GFP mouse tibia (3PM at 1330 nm). Scale bar = 50 µm.

**Video Captions**

**Video 1**: z-stack acquired in the intact tibia explanted from a C57/Bl6 mouse by three-photon-imaging at 1650 nm, 2 MHz rate repetition rate. The 3D image shows THG signal in a tissue volume of 442x442x362 µm³, 1036x1036x362 voxel, covering both cortical bone and bone marrow. The z-stack was used to determine the depth dependent spatial resolution in bone cortex and marrow, in the intact tibia (Fig. 3). Scale bar = 100µm.

**Video 2**: z-stack acquired in the intact tibia of a *Cdh5:tdTomato x Histone:GFP* mouse by *in vivo* three-photon imaging at 1650 nm excitation, with 1 MHz, 2 MHz, 3 MHz or 4 MHz repetition rate. 3D images show tdTomato fluorescence in the membranes of endothelial cells forming the blood vessel walls, within a tissue volume of 400x400x500 µm³, 751x751x126 voxel, the top layer laying in 70 µm tissue depth in the bone cortex. The data were used to determine the depth dependent signal-to-noise-ratio in the bone marrow (Fig. 4a). Scale bar = 100µm.

**Video 3**: *In vivo* 2D time lapse imaging (400x400 µm², 518x518 pixel) in the tibia marrow of a *Prx1:tdRFP* mouse performed at 1650 nm, 3 MHz, over 3 minutes, every second (Fig. 4c), followed by time lapse 2D imaging of a close up (150x150 µm², 257x257 pixel) in the same region, over 1 minute, every 0.5 s. Laser pulse energy, average laser power and imaging area location in tibia are indicated in **Table II**. tdRFP fluorescence (magenta) highlights the stromal compartment in the bone marrow and osteocytes in cortical bone, THG signal (green) reveals tissue architecture in both bone cortex and marrow. THG signal of erythrocytes highlights blood flow within the vascular system in a label-free manner. Scale bar = 50 µm.

**Video 4**: *In vivo* 2D time lapse imaging (400x400 µm², 518x518 pixel) in the tibia marrow of a *Prx1:tdRFP* mouse performed at 1650 nm, 4 MHz, over 3 minutes, every second (Fig. 4d), followed by time lapse 2D imaging of a close up (150x150 µm², 257x257 pixel) in the same region, over 40 s, every 0.5 s. Laser pulse energy, average laser power and imaging area location in tibia are indicated in **Table II**. tdRFP fluorescence is shown in magenta, THG signal in green. Blood flow within the vascular system is visualized by the THG signal of erythrocytes. Scale bar = 50 µm.

**Video 5**: *In vivo* 3D time lapse imaging (400x400x30 µm², 518x518x11 voxel) in the tibia marrow of a *CD19:tdRFP* mouse performed by 2PM at 1100 nm, 80 MHz, over 60 minutes, every 30 s, followed by 3PM at 1650 nm, 3 MHz, again over 60 minutes, every 30 s (Fig. 5). The tibia cortex was mechanically thinned to allow 2PM imaging in the bone marrow. Laser pulse energy, average laser power and imaging area location in tibia are indicated in **Table III**. tdRFP fluorescence (magenta) highlights B lineage cells in the bone marrow. Scale bar = 100 µm.

**Video 6**: *In vivo* 3D time lapse imaging (400x400x30 µm², 518x518x11 voxel) in the tibia marrow of a *CD19:tdRFP* mouse performed by 3PM at 1650 nm, 3 MHz, over 60 minutes, every 30 s, followed by 120 minutes, every 120 s (Fig. 6). Laser pulse energy, average laser power and imaging area location in the intact tibia, with 200 µm thick cortical bone, are indicated in **Table III**. tdRFP fluorescence (magenta) highlights B lineage cells in the bone marrow. Scale bar = 100 µm.

**Video 7**: *In vivo* 3D time lapse imaging (400x400x30 µm², 518x518x11 voxel) in the tibia marrow of a *CD19:tdRFP* mouse performed by 3PM at 1650 nm, 4 MHz, over 120 minutes, every 120 s (Suppl. Fig. 8). Laser pulse energy, average laser power and imaging area location in the intact tibia are indicated in **Table III**. tdRFP fluorescence (magenta) highlights B lineage cells in the bone marrow. Scale bar = 100 µm.

**Video 8**: *In vivo* 3D time lapse imaging (400x400x30 µm², 518x518x11 voxel) of label-free THG signal in the intact mouse tibia performed by 3PM at 1650 nm, 4 MHz, over 120 minutes, every 120 s. Laser pulse energy, average laser power and imaging area location in the intact tibia are indicated in **Table III**. THG signals (green) show tissue architecture and strain in bone cortex and bone marrow. During the video, the focus dived from the bone cortex, into the bone marrow, to highlight strain dynamics in tissues of different stiffness. Scale bar = 100 µm.

**Video 9**: *In vivo* 3D time lapse imaging (400x400x30 µm², 518x518x11 voxel) in the tibia marrow of a *CD19:tdRFP* mouse performed by 3PM at 1650 nm, 3 MHz, over 60 minutes, every 30 s (Fig. 7). Laser pulse energy, average laser power and imaging area location in the intact tibia are indicated in **Table III**. tdRFP fluorescence (magenta) highlights B lineage cells in the bone marrow and $THG^{hi}$ signal (green) indicates cells in the bone marrow with abundant endoplasmic reticulum. Scale bar = 100 µm.
